# *Limosilactobacillus reuteri* promotes the expression and secretion of enteroendocrine- and enterocyte-derived hormones

**DOI:** 10.1101/2024.08.30.610555

**Authors:** Sara C. Di Rienzi, Heather A. Danhof, Micah D. Forshee, Ari Roberts, Robert A. Britton

## Abstract

Observations that intestinal microbes can beneficially impact host physiology have prompted investigations into the therapeutic usage of such microbes in a range of diseases. For example, the human intestinal microbe *Limosilactobacillus reuteri* strains ATCC PTA 6475 and DSM 17938 are being considered for use for intestinal ailments including colic, infection, and inflammation as well as non-intestinal ailments including osteoporosis, wound healing, and autism spectrum disorder. While many of their beneficial properties are attributed to suppressing inflammatory responses in the gut, we postulated that *L. reuteri* may also regulate hormones of the gastrointestinal tract to affect physiology within and outside of the gut. To determine if *L. reuteri* secreted factors impact the secretion of enteric hormones, we treated an engineered jejunal organoid line, *NGN3*-HIO, which can be induced to be enriched in enteroendocrine cells, with *L. reuteri* 6475 or 17938 conditioned medium and performed transcriptomics. Our data suggest that these *L. reuteri* strains affect the transcription of many gut hormones, including vasopressin and luteinizing hormone subunit beta, which have not been previously recognized as being produced in the gut epithelium. Moreover, we find that these hormones appear to be produced in enterocytes, in contrast to canonical gut hormones which are produced in enteroendocrine cells. Finally, we show that *L. reuteri* conditioned media promotes the secretion of several enteric hormones including serotonin, GIP, PYY, vasopressin, and luteinizing hormone subunit beta. These results support *L. reuteri* affecting host physiology through intestinal hormone secretion, thereby expanding our understanding of the mechanistic actions of this microbe.

## Introduction

The use of commensal microbes in the treatment of disease has the potential to herald in a new era of microbial-based therapeutics. The human associated *Limosilactobacillus reuteri* is one such microbe considered for development as a therapeutic: it has been shown to improve symptoms of infant colic^1^, osteoporosis^2^, and inflammatory diseases^3–6^, and is being considered for its role in alleviating asocial behavior associated with autism spectrum disorder^7–11^. How *L. reuteri* mediates these effects is not fully understood. Moreover, several different *L. reuteri* strains are currently in use, highlighting the importance of studying strain variation in understanding therapeutic efficacy.

Two of the commonly employed strains that are currently marketed as probiotics are *L. reuteri* ATCC PTA 6475 and *L. reuteri* DSM 17938. While both were originally derived from human breast milk, these strains are phylogenetically and functionally distinct. *L. reuteri* 6475, belongs to *L. reuteri* clade II, while *L. reuteri* 17938 (derived from strain ATCC 55730^12^) belongs to *L. reuteri* clade VI^13^. *L. reuteri* 17938 (or its parent *L. reuteri* 55730) has been demonstrated to reduce infant colic^1^, assist in feeding tolerance in preterm infants^14^, improve intestinal motility in preterm^14^ and term infants^15^, and improve cytokine ratios in children with apoptotic dermatitis^16^. *L reuteri* 6475 has been shown to have potential in relieving inflammatory conditions through TNF suppression, which may be linked to its capacity to reduce osteoporosis^2,17–23^. *L. reuteri* 6475 has also been demonstrated efficacious in promoting wound healing^24,25^, restoring normal social behavior in mouse models of autism^7–9,11^ (which *L. reuteri* 17938 has been shown unable to do so in mice), and improving male reproductive health in mice^26^. These two strains are similar in their ability to produce the antimicrobial reuterin and the vitamins pseudo B12 and B9 (folate)^27^ and to produce proteins for host mucus adherence^28^. *L. reuteri* 6475 can also produce histamine while *L. reuteri* 17938 cannot^13^. This histamine production is implied in *L. reuteri* 6475’s suppression of the inflammatory signal tumor necrosis factor (TNF)^27^. *L. reuteri* 17938 has also been demonstrated to liberate adenosine from AMP, which may be involved in its function to reduce autoimmunity in Treg deficiency disorders by enhancing CD73^+^CD8^+^T cells^29^.

While many of *L. reuteri*’s functions are thought to be due to interactions with immune cells, *L. reuteri* itself or its secreted products has the capacity to influence host physiology through a wide range of cell types. Particularly in the small intestine, where the mucus layer is thin, *L. reuteri* may have ample opportunities to interact with the host epithelial cells. Given, the diverse roles of *L. reuteri* in gut motility, on inflammatory processes, and on the gut-brain axis led us to consider whether some of *L. reuteri*’s interactions with the host are mediated through enteroendocrine cells.

Enteroendocrine cells are secretory cells in the intestine specialized for the secretion of hormones. Enteroendocrine cells sense nutrients like sugars, peptides, and fatty acids in the intestinal lumen through G-protein coupled receptors and utilize ion (sodium, hydrogen, calcium) transporters to bring nutrients into the cell^30^. On apical entry or basolateral exit from enteroendocrine cells, these nutrients can trigger hormone receptors and lead to the release of hormones from the apical or basolateral side of the cell^31^.

Enteroendocrine cells also respond to microbial stimulus through toll-like receptors to release cytokines and subsequently affect inflammatory responses^30^. As well, released gut hormones can directly and indirectly influence pro- and anti-inflammatory immune cell populations through a variety of mechanisms^30^. Finally, enteroendocrine cells and a few specific hormones are associated with the integrity of the intestinal barrier^30^.

Enteroendocrine cells, however, comprise ∼1% of gut epithelial cells, thereby making study of these cells difficult *in vivo* and in non-transformed tissue lines. To overcome this limitation, we recently developed a human enteroendocrine-enriched jejunal organoid line^32^. Through induction of the developmental regulator of enteroendocrine cells, *NGN3*, we can increase the number of enteroendocrine cells to ∼40% in this adult cell stem derived human jejunal organoid line at the expense of enterocytes^32^.

Here, we utilized these *NGN3* human intestinal organoids (HIOs) to characterize how *L. reuteri* secreted products impact enteroendocrine cells. By performing RNA-Seq on uninduced organoids and induced, enteroendocrine-enriched organoids, we observe that *L. reuteri* affects the transcription of genes involved in hormone secretion, nutrient sensing, cell adhesion, mucus production, immune/stress response, and cell fate. Among the impacted hormones are enterocyte-derived hormones, not previously characterized in the intestinal epithelium. For several of the impacted hormones, we additionally demonstrate that *L. reuteri* promotes the secretion of these hormones from HIOs or from *ex vivo* human intestinal tissue. In general, we observe similar effects of *L. reuteri* strains 6475 and 17938 on epithelial cells but with *L. reuteri* 6475 having a greater magnitude of effect on transcription. These results suggest specific mechanisms by which *L. reuteri* mediates its beneficial effects with a magnified look at how *L. reuteri* interacts with enteric hormones.

## Methods

### Preparation of bacterial conditioned media

*L. reuteri* strains ATCC PTA 6475 and DSM 17938 were provided by BioGaia (Sweden). A single colony of *L. reuteri* 6475 or 17938 from an MRS agar plate was inoculated into 10 mL of MRS broth and incubated in a tightly closed conical tube in a 37°C water bath or incubator. After 15 hours of incubation, the *L. reuteri* culture was diluted to an OD_600_ of 0.1 into 25 to 40 mL of pre-warmed LDM4^6^ and placed into a 37°C water bath to incubate until reaching an OD_600_ of 0.5-0.6. Next, cells were pelleted by centrifugation and the resulting supernatant was transferred to a new conical tube. The pH of the supernatant was measured by applying 2 µL of the supernatant onto pH paper (range 6.0 – 8.0, Fisherbrand, Pittsburgh, PA, USA) and adjusted to 7.0 using 10 M sodium hydroxide solution. Neutralized conditioned media and LDM4 media control were filter sterilized (0.22µm PVDF membrane, Steriflip, EMD Millipore, Burlington, MA), aliquoted, frozen at -80°C overnight, and then lyophilized. Lyophilized conditioned media were stored at -20°C until use.

### Propagation of organoids and organoid media

J2 *NGN3* organoids were propagated in 3D in CMGF+ media^32^ + 10 µmol Y-27632 Rock inhibitor + 200 µg/ml geneticin as previously described^33^. *NGN3*-HIOs were then seeded onto 24-well transwells and differentiated in the presence of differentiation media^32^ with (induced) or without (uninduced) 1 µg/ml doxycycline.

### Transwell assay

For use on organoids, lyophilized conditioned media were resuspended in an equal volume of organoid differentiation media. The existing differentiation media on the apical side of the transwells were removed and replaced with 100 µL differentiation media supplemented with lyophilized conditioned media or media control. Transwells were incubated for 3 hours at 37°C with 5% CO_2_. Following, apical and basolateral supernatants were removed and stored at -20°C in a 96 well plate to be used later in a hormone secretion assay. The transwell membrane was removed from the support surface and placed in TRIzol solution (Invitrogen, Waltham, MA, USA). Following a chloroform extraction, the aqueous phase containing total RNA was immediately extracted using a Qiagen RNeasy kit (Qiagen, Germantown, MD, USA).

### RNA-Seq

Paired-end Illumina sequencing libraries were prepared by Novogene (Sacramento, CA, USA). Briefly, total RNA was enriched for Eukaryote mRNA. mRNA was fragmented to an average insert size of 250 to 300 bp, and cDNA was prepared using the standard NEB library construction method. The library was 150 bp paired-end sequenced on a NovaSeq 6000. Basecalling was performed using CASAVA v1.8^34^.

Reads were filtered as follows: reads containing adaptors were removed, reads with more than 10% N reads were removed, and reads where > 50% of the bases have Qscore <= 5 were removed.

Sequenced reads were aligned to the human genome hg19 using Star (v2.5)^35^ using the Maximal Mappable Prefix for junction reads and with mismatch = 2. Read counts per gene were tabulated with HTSeq v0.6.1^36^. The gene count table provided by Novogene was further processed using a pipeline derived from iDEP version 0.82^37^. Genes were filtered to keep those with at least 1 count per million in 5 samples, thereby retaining 15,369 genes.

For multidimensional scaling, rlog transformed data were visualized using a t-distribution to estimate the hypothetical spread of the data. The contribution of induction and *L. reuteri* treatment to the variation in data were modeled using a permutational multivariate analysis of variance (PERMANOVA) of the form: Euclidean distance matrix ∼ induction + treatment + induction * treatment using the adnois function in vegan (v2.5-5)^38^.

For correlation analyses, rlog-transformed values were used. Lowly expressed genes belonging to the bottom quartile were removed. Correlations among samples were computed using a Pearson correlation. Correlations were visualized using the ComplexHeatmap package (v2.3.1)^39^, with rows and columns clustered by a Euclidean distance metric and using complete linkage clustering for both. Within and between sample distances were plotted using the ggboxplot function in ggpubr (v0.2.4)^40^. Significance among distances was calculated by a t-test with a multiple testing correction using Holm’s method^41^.

Difference between means (circle size) and adjusted p-values (circle color) were visualized as a correlogram using ComplexHeatmap package (v2.3.1)^39^.

For identification of differentially expressed genes, gene counts were modeled as genecount ∼ treatment- induction + organoid_batch in DESeq2^42^ V1.22.2 using a Wald test with p values corrected using the Benjamini-Hochberg procedure^43^ with an FDR cutoff of 0.1 and a fold change cutoff of 2. DESeq2 models the underlying variation using a negative binomial distribution. LDM4 (media alone) and uninduced (not enteroendocrine enriched) were used as reference levels.

### Functional analyses

Ensembl IDs release 95 were converted to Ensembl IDs release 98 before analyzing for statistical enrichment of gene functions using the Ensembl ID converter^44^. Annotations for PANTHER GO-Slim Biological Process, PANTHER GO-Slim Molecular Function, PANTHER GO-Slim Cellular Component, PANTHER Protein Class, Panther Pathways, and Reactome^45,46^, were performed in PANTHER^47^, using a binomial test, and a false discovery cutoff of 0.05. Genes belonging to enriched (not depleted) functional categories defined by PANTHER^47^ were searched in GeneCards^48^ and annotated into one of the following broad groups: Cell fate/growth, Hormone secretion, Immune response, Membrane component, Mucus, Nutrient metabolism/response, Signaling, or Metal/stress response. Enrichments of these groups within Kmeans determined clusters (see below for heatmap visualization) were determined using a hypergeometric distribution, and all p-values across groups and clusters were corrected *en masse* using the Benjamini-Hochberg^43^ method, whereby FDR values less than 0.1 were considered significant.

### Data visualization

For multidimensional scaling, clustering, and heatmap visualization, read counts were transformed using the rlog function from DESeq2 V1.22.2^42^. For displaying the gene expression data as a heatmap, the rlog transformed data were batch corrected using the removeBatchEffect command in the limma package^49^ and the data were centered and scaled using the scale function in base R^50^. Heatmaps were visualized using the ComplexHeatmap package^39^, with rows (genes) clustered with the Pearson distance metric and columns (samples) clustered with the Euclidean distance metric, using complete linkage clustering for both. The number of clusters to group the displayed genes was determined using the Kmeans function in base R^50^, with visualization of the total sum of squares as an elbow plot and average silhouettes in a silhouette plot. The number of clusters to group the samples (columns) was selected solely for enhancing visualization. For barplots of individual gene expression values, read counts were transformed using the GeTMM method^51^ and converted to counts per million using calcNormFactors and cpm commands in edgeR^52^. Displayed log_2_ fold changes were derived from DESeq2 modeled data. In this method, the log fold changes are shrunken to prevent overestimation of fold changes for genes with low counts and/or high dispersion. Enterocyte and enteroendocrine cell markers were referenced from Haber and colleagues^53^.

### Gene annotations

Annotations for select hormone-related genes were taken from GeneCards^48^ (www.genecards.org) and from the literature: AGT^54,55^, ARHGEF25^56^, CCK^57–59^, GAST^57,60,61^, GHRL and GHRLOS^62–64^, GIP^60^, MLN^57,60^, NPW^65–67^, NPY^68,69^, SST^70–72^, DRD1^73^, NRG4^74–76^, NTSR1^60,77^, TAC3^68,78,79^, AVP^80–85^, C1QTNF12^86,87^, LHB^88,89^, NTS^60,77^, OXT^8,84,85,90,91^, SCTR^92^, PAQR5^93^, P2RY1^94^, RARB^95^. Annotations for select immune and stress response genes were taken from GeneCards^48^ (www.genecards.org).

### Human tissue

Human intestinal tissue was acquired from the organ donation group LifeGift within the Texas Medical Center. All organ donors were adults not presenting with any known gastrointestinal disease, surgery, or trauma. Individuals positive for hepatitis B or C, HIV, or COVID were excluded. Tissue was delivered to lab within ∼4 hours of the patient initiating organ harvest and within ∼1 hour of harvest of the gastrointestinal tract.

### Hormone secretion

To measure secreted hormones from the treated organoids, supernatants from the apical (or basolateral, where noted) side of the transwells were assessed using the Luminex MILLIPLEX Human Metabolic Hormone kit (EMD Millipore, USA) or using a serotonin ELISA (SER39-K01, Eagle Biosciences, USA). For measuring hormones secreted from whole human tissue, approximately 2 cm by 2 to 3 cm pieces of human tissue were incubated in 5 mLs of *L. reuteri* conditioned media or media control in 6 well plates for 3 hours at 5% CO_2_. AVP was measured with the Arg8-Vasopressin ELISA kit (ADI-900-017A, Enzo, USA), LHB with the Luteinizing Hormone (hLH) ELISA Assay kit (HLH31-K01, Eagle Biosciences, USA), adipolin with the Human CTRP12 ELISA kit (SK00392-06, Aviscera Bioscience, USA), and kisspeptin with the Human Kisspeptin ELISA kit (ab288589, Abcam, USA). For organoids, statistical significance was determined using a one-way ANOVA followed by a Dunnett’s test with the LDM4 treatment used as the control. For human tissue, data were modeled with linear mixed models with the human patient included as a random variable using the lmer function of the lme4^96^ package with REML = FALSE and the control optimizer = “bobyqa”. Following, statistical significance was determined using the emmeans function^97^ with a Benjamini-Hochberg multiple testing correction.

### Single cell RNA-Sequencing analysis

Single cell RNA-Sequencing (scRNA-Seq) analysis of the Human Gut Atlas (https://www.gutcellatlas.org/, adult epithelium, jejunum) was performed as previously described^33^. Briefly, scRNA-Seq data from the adult jejunum were analyzed using the Seurat package in R (v 5.0.3). After data normalization, data clustering, and UMAP generation, genes of interest were plotted using the FeaturePlot function.

### Immunofluorescence

Adipolin was visualized on human jejunal tissue from organ donors as previously described^33^ using the antibody (NBP1-90700, Novus Biologicals) diluted to between 1:20 to 1:50 and detected with Rhodamine Red™-X (AB_2338028, Jackson ImmunoResearch) diluted to 1:200. An E-caderin conjugated antibody (1:50, 560062, BD Pharmingen) and DAPI (1:10, NucBlue™ Fixed Cell Stain ReadyProbes™ reagent, R37606, Invitrogen) were applied simultaneously with Rhodamine Red™-X.

### Data availability

RNA-Seq reads are available at NCBI GEO at https://www.ncbi.nlm.nih.gov/geo/ accession number GSE138350 and GSE268681. Scripts for plots and data are available at https://github.com/sdirienzi/Lreuteri_HIORNASeq. An interactive ShinyApp displaying the RNASeq data can be found here: https://sdirienzi.shinyapps.io/LreuHIORNASeq/.

## Results

### *NGN3*-HIOs facilitate study of *L. reuteri*’s interactions with the enteroendocrine system

To determine how *L. reuteri* strains 6475 and 17938 affect the intestinal epithelium, we designed an RNA-Seq experiment using human intestinal organoids (HIOs) treated with pH neutralized conditioned media produced by these strains in log phase (**Figure 1A**). The media thereby represent any products released by the *L. reuteri* strains into their growth media. The specific HIOs we utilized originated from adult jejunal stem cells and have been engineered for the inducible expression of the transcription factor *NGN3*. *NGN3* induction results in HIOs enriched in enteroendocrine cells with a decrease in the relative abundance of enterocytes^32^. With this *NGN3*-HIO line we can measure the effects of the *L. reuteri* strains on induced *NGN3*-HIOs enriched in enteroendocrine cells and on uninduced *NGN3*-HIOs largely comprised of enterocytes.

**Figure 1.**
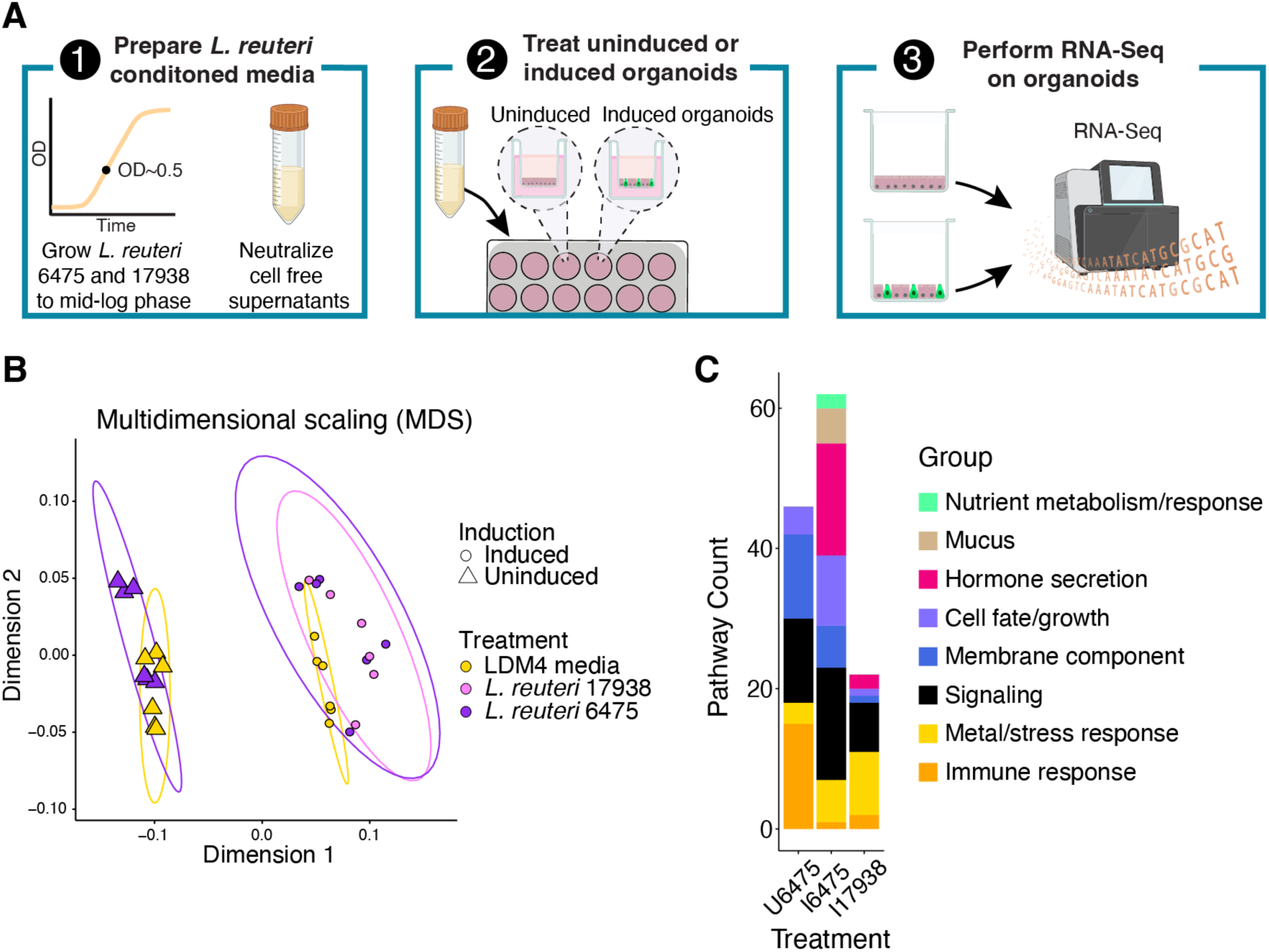
Induced and uninduced *NGN3*-HIOs differentially respond to *L. reuteri* treatment. **A)** Overview of RNA-Seq experiment. First, *L. reuteri* conditioned media was prepared by growing *L. reuteri* 6475 and 17938 in LDM4 to mid-log phase. The bacterial cells were spun out, the resulting conditioned media brought to neutral pH, and then filtered through a 0.22 µm filter. The conditioned media were then lyophilized and resuspended in HIO differentiation media. These treatments were then placed into uninduced or induced *NGN3*-HIOs in transwells for three hours. Third, the organoid cells were harvested, and isolated RNA was sent for RNA-Seq. Created with BioRender.com. **B)** Principal coordinate analysis of transcriptomic data from *NGN3*-HIOs induced or not induced and treated with *L. reuteri* 6475, 17938, or LDM4 media control. Ellipses for illustration purposes are modeled from the data following a t-distribution. **C)** Enriched functional categories of differentially expressed genes in *L. reuteri* treatments over media alone. U6475 is *L. reuteri* 6475 vs media control in uninduced *NGN3*-HIOs. I6475 is *L. reuteri* 6475 vs media control in induced *NGN3*-HIOs. I17938 is *L. reuteri* 17938 vs media control in induced *NGN3*-HIOs. Some functional groups are listed as belonging to two categories (see Supplemental Table 3 for further details).

We tested *L. reuteri* 6475 on uninduced *NGN3*-HIOs (∼90% enterocytes, <2% enteroendocrine cells)^32^ and *L. reuteri* 6475 and 17938 on enteroendocrine-enriched (induced) *NGN3*-HIOs (∼50% enterocytes, ∼40% enteroendocrine cells)^32^. *L. reuteri* 17938 was not tested on uninduced HIOs. RNA-Seq of the organoids produced an average of 16.1 million reads per library (**Table 1**, **Supplemental Table 1**). To confirm that induced *NGN3*-HIOs were enriched in enteroendocrine cells and depleted in enterocytes compared to uninduced *NGN3*-HIOs, we checked the expression level of known enterocyte and enteroendocrine cell markers^53^ (**Figures S1** and **S2**). The expression levels of genes followed the expected patterns with enterocyte markers being downregulated and enteroendocrine markers increasing with *NGN3* induction (**Figures S1** and **S2**).

**Table 1:** Summary of RNA-Seq libraries. Read counts shown are post filtering and alignment to the human genome (See Methods). See Supplemental Table 1 for further details.

| HIO type | Treatment | Number of HIO experiments | Number of replicate HIOs within an experiment | Total number of RNA-Seq libraries | Average read count |
| --- | --- | --- | --- | --- | --- |
| Uninduced | LDM4 | 2 | 3 | 6 | 18968834.33 |
| Uninduced | *L. reuteri* 6475 | 2 | 3 | 6 | 15465905.00 |
| Enteroendocrine-enriched | LDM4 | 2 | 3 | 6 | 15972039.33 |
| Enteroendocrine-enriched | *L. reuteri* 6475 | 2 | 3 | 6 | 14287207.00 |
| Enteroendocrine-enriched | *L. reuteri* 17938 | 2 | 3 | 6 | 15720129.33 |

To globally assess whether the HIOs were impacted by the *L. reuteri* conditioned media, we performed an unsupervised analysis using dimensionality reduction with multidimensional scaling (MDS) produced from a Euclidean distance matrix of the gene expression data. As expected, the MDS plot illustrated that the data could be separated in dimension 1 by whether the HIOs were induced for *NGN3* expression or not, indicating *NGN3* induction was likely the greatest contributor to the variation in global gene expression (**Figure 1B**). To quantify the contribution of induction as well as the contributions of *L. reuteri* treatments and biological replication, we performed a PERMANOVA on the Euclidean distance matrix. Our PERMANOVA model reported that *NGN3* induction explains 70.8% of the variation (pseudo-F = 120.763, p = 0.001), biological replication 9.2% of the variation (15.685, p = 0.001), *L. reuteri* treatment 4.4% of the variation (pseudo-F = 3.768; p = 0.011), and that the interaction of treatment and induction was not significant (1.5% of the variation; pseudo-F = 2.578; p = 0.082). Similar results were obtained using the Jaccard similarity index. These results indicated that most of the variation in data resulted from *NGN3* induction, and that the addition of *L. reuteri* 6475 or 17938 had a relatively smaller but still significant effect on HIO gene expression.

To gain further insight into the variation in gene expression in our data, we investigated gene expression correlations among pairwise comparisons of samples. We observed that induced HIOs treated with either *L. reuteri* strain were significantly less correlated from *L. reuteri* 6475 vs media control on uninduced HIOs (**Supplemental Figure 3A**, **Supplemental Figure 3B**). We also observed that the correlations between induced HIOs treated with *L. reuteri* 6475 vs their media controls compared to those treated with *L. reuteri* 17938 vs their media controls were similar (p=0.09), although the mean correlation for induced HIOs treated with *L. reuteri* 6475 vs their media controls was lower (**Supplemental Figure 3A, B**). Together, these results further support that both *L. reuteri* strains had a significant effect on HIO gene expression when the HIOs were induced and suggest that the *L. reuteri* strains similarly affected gene expression.

### *L. reuteri* strains 6475 and 17938 impact the expression of hormone, nutrient, mucus, metal/stress response, and immune-related genes in native and/or enteroendocrine-enriched HIOs

We next sought to determine the genes impacted by *L. reuteri* strains 6475 and 17938 in the induced and uninduced *NGN3* HIOs. We identified differentially expressed genes (DEGs) between these two strains and across the induction state of the HIOs. Specifically, we compared the effect of *L. reuteri* 6475 in the uninduced and induced states compared to their media controls and *L. reuteri* 17938 in the induced state to its media control. We find a similar number of genes impacted by *L. reuteri* 6475 in induced and uninduced HIOs, but fewer DEGs by *L. reuteri* 17938 in induced HIOs (**Table 2**, **Supplemental Table 2**). While at first glance, this may suggest *L. reuteri* 6475 affects HIO differently than 17938, only 12 genes were differentially expressed between *L. reuteri* 6475 and 17938 in induced HIOs (**Table 2; Supplemental Figure 3C**). On investigating the gene expression data, we observed that *L. reuteri* 17938 largely affects gene expression in the same direction as 6475, but that the fold change in gene expression for 17938 failed to pass our significance thresholds. These results reinforce the results of our correlation analysis (**Supplemental Figure 3**) suggesting that though *L. reuteri* 6475 had a more potent effect on transcriptional change in our induced HIOs in this experimental setup, the two strains had largely similar effects on gene transcription.

**Table 2:**
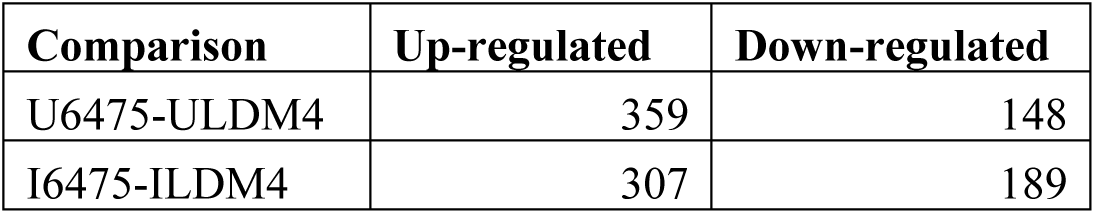
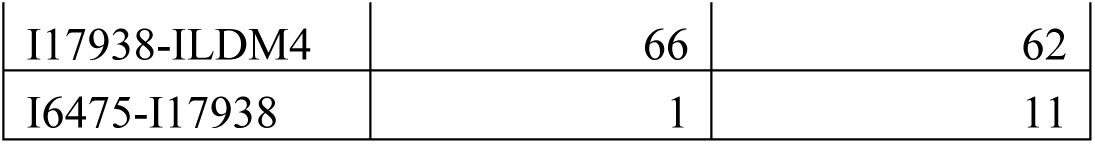
Summary of genes differentially regulated between *L. reuteri* strains 6475 and 17938 in induced and uninduced HIOs. . Groups are labeled as “U” for uninduced, “I” for induced, “6475” for treatment with *L. reuteri* 6475 conditioned medium, “17938” for treatment with *L. reuteri* 17938 conditioned medium, and “LDM4” for treatment with bacterial growth medium.

To determine how these transcriptional changes might functionally affect the HIOs, we looked for functional enrichments in the DEGs. Using the PANTHER classification system^47^ and the Reactome annotated pathways^45,46^, we identified enriched functional annotations within the sets of DEGs. Broadly across all datasets, the *L. reuteri* DEGs were enriched in functions regarding response to the environment. These functions included those for nutrient, stress, metal, and immune response, cell fate/growth, membrane components, and signal transduction (**Figure 1C & Supplemental Table 3**). As anticipated, the induced HIOs treated with either *L. reuteri* strain were also enriched in genes for hormone secretion (**Figure 1C & Supplemental Table 3**). The induced cells treated with *L. reuteri* 6475 were additionally enriched for genes relating to mucus.

To further investigate and understand the DEGs and their regulation, we annotated these genes within the functional groups and looked for similar expression patterns and functions (**Supplemental Table 3**). We were able to group the genes into 8 groups using Kmeans clustering (**Supplemental Figure 4**). These clusters represent genes with similar transcriptional responses to induction and the presence of *L. reuteri* and therefore may share similar regulatory mechanisms. For instance, genes within a cluster may share a transcription factor or be localized within the same cell type. As cell types within the small intestine have partially non-overlapping functions^98^, this scenario would promote clusters being enriched in one or two closely related functions.

Indeed, we observed this to be the case (**Table 3**): clusters were either enriched in one or two related functions or were not enriched in any function. Clusters 1 and 5 were enriched in genes involved in hormone secretion; cluster 2 in cell adhesion; cluster 3 in stress/immune response; cluster 6 in nutrient response; and clusters 7 and 8 in mucus genes. Therefore, the clusters generated by our heatmap are consistent with gene clusters of related functionalities, perhaps from genes expressed in the same or similar cell type.

**Table 3:** Functional enrichments within clusters of similarly expressed DEGs. Clusters are listed as in Supplemental Figure 4. Columns U6475-ULDM4 through ILDM4-ULDM4 summarize whether genes within that cluster are predominantly up (+) or down (-) regulated for the given comparison. FDR corrected significance values for functional groups: *, q<0.1, **, q<0.01, ***, q<0.001.

| Cluster | U6475-ULMD4 | I6475-ILDM4 | I17938-ILDM4 | I6475-U6475 | ILDM4-ULDM4 | Cell<br>adhesion | Cell fate | Hormone<br>secretion | Mucus | Nutrient<br>response | Signaling | Stress/<br>Immune<br>response |
| --- | --- | --- | --- | --- | --- | --- | --- | --- | --- | --- | --- | --- |
| 1 | - | + |  | + | + | 0.454 | 0.793 | **0.094\*** | NA | 0.542 | **0.001\*\*** | 1 |
| 2 | - | - | - | + | + | **0.064\*** | 0.879 | 0.879 | 0.231 | 0.542 | 0.542 | 0.454 |
| 3 | + | + | + | + | + | NA | 0.82 | 0.454 | NA | 0.82 | 0.879 | **0.021\*** |
| 4 | + |  |  | - |  | NA | 0.106 | NA | NA | 0.542 | 0.399 | 0.655 |
| 5 | + | + | + | - | - | NA | 0.542 | **0.064\*** | NA | 0.283 | NA | 0.454 |
| 6 | - |  |  | - | - | NA | 0.454 | 0.769 | NA | **0.037\*** | NA | 0.399 |
| 7 |  | - | - | - | - | 0.231 | 0.454 | 0.879 | **0.064\*** | 0.879 | 0.542 | 0.542 |
| 8 |  | - |  |  |  | NA | 0.283 | 0.769 | **0.086\*** | NA | 0.772 | 0.399 |

### *L. reuteri* impacts on immune and stress response

To see if our data are consistent with known functions of *L. reuteri* on the intestinal epithelium, we first investigated the immune and stress response DEGs. We observed many immune-related genes were downregulated and a few metal and stress response genes were upregulated by *L. reuteri* 6475 (**Supplemental Figure 5A**). Tumor necrosis factor (TNF), which *L. reuteri* 6475 has been previously observed to downregulate^17^ and suppress^3^, was not expressed in our HIOs; however, TNFSF15, which is induced by TNF and activates NF-kappaB^99^, was decreased in induced HIOs treated with *L. reuteri* 6475. Consistent with *L. reuteri* 6475 mediated suppression of NF-kappaB and inflammatory responses^17^, several chemokines were downregulated by *L. reuteri* 6475: IL-8 (CXCL8), CCL2 (MCP-1), CXCL2, CX3CL1, and CXCL3. Secreted MCP-1, we observe, was also repressed by both *L. reuteri* strains only on induced *NGN3*-HIOs (**Supplemental Figure 5B**), suggesting a role of enteroendocrine cells in downregulating inflammatory responses. TLR4 which senses stimuli and upregulates inflammatory responses^100^ was also downregulated by *L. reuteri* 6475. *L. reuteri* 6475 additionally upregulated interleukin 18 binding protein (IL18BP), which is an inhibitor of the proinflammatory IL-18^101^. Defensin- 6, interferon epsilon, several metallothionein genes, and aquaporin-1 and -7, which respond to environment changes, were upregulated by *L. reuteri* 6475. These data are consistent with reports of *L. reuteri* 6475 having anti-inflammatory, immune modulatory, and stress response effects on the gut epithelium. *L. reuteri* 17938 had a less pronounced effect on immune and stress response genes. None of the chemokine or aquaporin genes were significantly impacted and only about half of the metallothioneins were differentially regulated in response to *L. reuteri* 17938. As mentioned previously, these results largely appear to be the result of *L. reuteri* 17938 impacting gene expression in the same direction but not the same magnitude as 6475 in our experiment.

### *L. reuteri* affects the transcription and secretion of enteroendocrine cell hormones

We next focused on clusters 1 and 5 for their enrichment of hormone genes (**Figure 2**). Cluster 1 appears as we would expect for canonical gut hormones derived from enteroendocrine cells: the genes in cluster 1 increased in expression with *NGN3* induction. These genes included those for the hormones angiotensinogen (AGT), cholecystokinin (CCK), gastrin (GAST), ghrelin (GHRL and GHRLOS), gastric inhibitory polypeptide aka glucose dependent insulinotropic polypeptide (GIP), motilin (MLN), neuropeptide W (NPW), neuropeptide Y (NPY), and somatostatin (SST). With the exception *AGT*, all genes were significantly upregulated by *L. reuteri* 6475. Only *GHRL* and *GHRLOS* were significantly upregulated by *L. reuteri* 17938.

**Figure 2:**
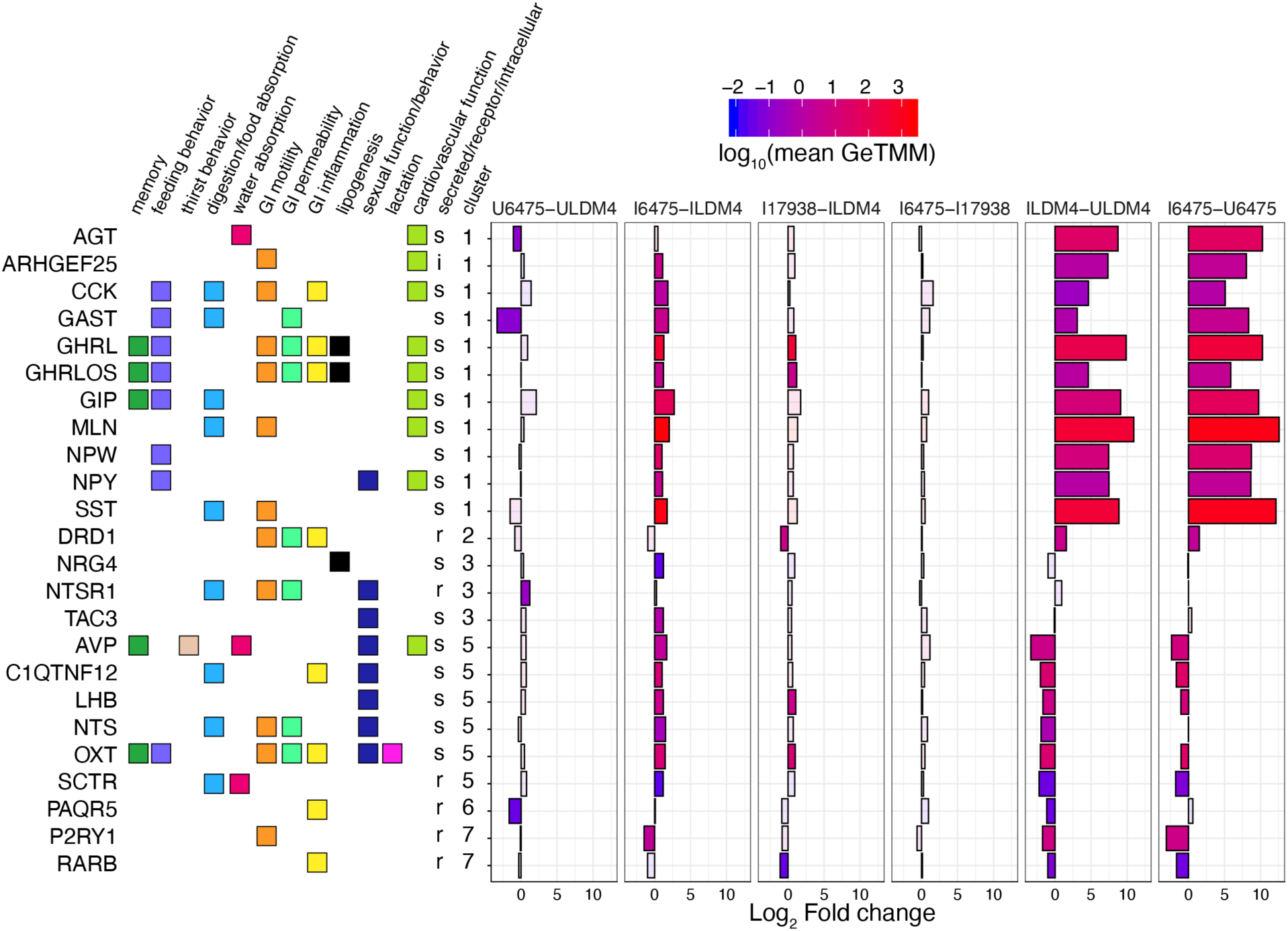
Hormone genes differentially expressed by *L. reuteri*. DEGs annotated as having hormonal function are shown. The genes are annotated with their function, whether they are secreted, a receptor, or intercellular, and what cluster they belong to as in Supplemental Figure S4. The graph shows the log_2_ fold change expression of the gene for the indicated comparison. The bars are colored using the log_10_ scaled mean GeTMM counts to illustrate how abundantly expressed the gene is. Transparent overlays are used on genes not differentially expressed for the given comparison. Comparisons shown: U6475-ULDM4, *L. reuteri* 6475 on uninduced HIOs compared to LDM4 media control; I6475-ILDM4, *L. reuteri* 6475 on induced HIOs compared to LDM4 media control; I17938-ILDM4, *L. reuteri* 17938 on induced HIOs compared to LDM4 media control; I6475-I17938 *L. reuteri* 6475 compared to *L. reuteri* 17938 on induced HIOs; ILDM4-ULDM4, LDM4 media control on induced versus uninduced HIOs; I6475-U6475, *L. reuteri* 6475 on induced versus uninduced HIOs. For each, positive fold changes indicate genes upregulated by the condition listed first.

To determine if some of these gene expression differences might lead to differences in hormone secretion, we tested the organoid supernatant that had been collected following the application of *L. reuteri* 6475 and 17938 conditioned media to uninduced and induced *NGN3*-HIOs. The harvested supernatants coming off the organoids were run on a Luminex panel consisting of metabolic-related hormones (see Methods) (**Figure 3A**). From this panel, we were able to obtain measurable values of amylin, C-peptide, ghrelin, GIP (total), pancreatic polypeptide (PP), and peptide YY (PYY) (**Figure 3B-G**). For amylin and PYY, both *L. reuteri* strains significantly increased secretion of these hormones from induced *NGN3*-HIOs (**Figure 3B, G**). The secretion of GIP was enhanced significantly (at p<0.05) by *L. reuteri* 17938 and C-peptide secretion was significantly promoted by *L. reuteri* 6475; although for both hormones, the other *L. reuteri* strain promoted secretion at p<0.1 (**Figure 3C, E**). PYY, who secretion was promoted, was not transcriptionally upregulated by either *L. reuteri* strain. PP (*PPY*), amylin (*IAPP*), and insulin (*INS*) gene counts were below the limit of detection in the RNA-Seq data.

**Figure 3.**
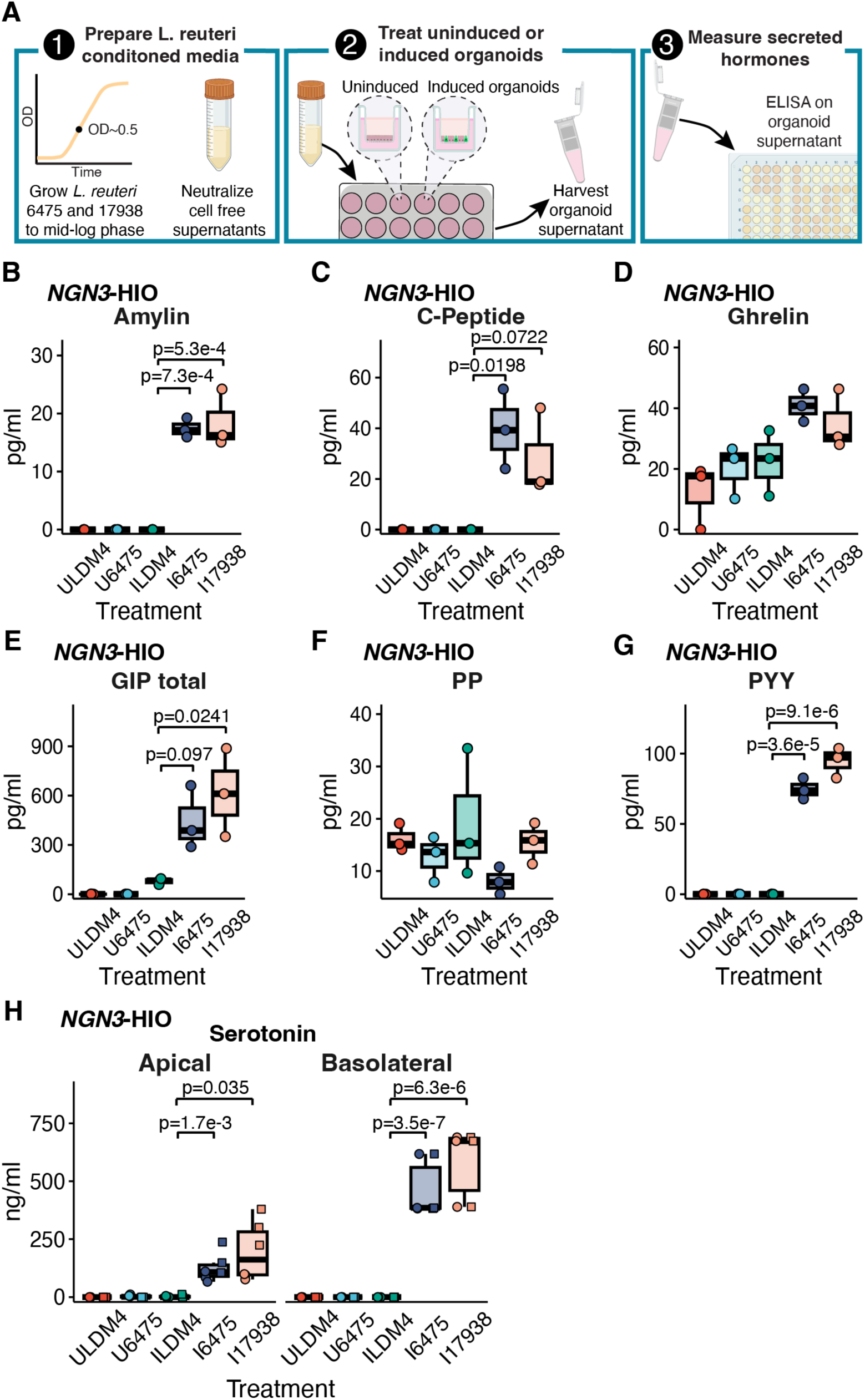
*L. reuteri* promotes the secretion of known enteroendocrine-derived intestinal hormones. **A**) 1) In order to measure the release of intestinal hormones from human intestinal organoids (HIO), *L. reuteri* conditioned media is generated from mid-log phase cultures of *L. reuteri*. These cultures are pH neutralized and rendered cell-free. 2) *L. reuteri* conditioned media is then placed onto *NGN3*-HIOs plated on transwells that are differentiated but not induced for *NGN3* or induced for *NGN3*. 3) Following an incubation on the HIOs, the supernatant is collected and secreted hormones are measured by ELISA or Luminex assay. Created with BioRender.com. Secreted amylin (**B**), C-peptide (**C**), ghrelin (**D**), GIP (**E**), PP (**F**), and PYY (**G**) measured from uninduced and induced *NGN3*-HIOs in response to *L. reuteri* 6475 or 17983 conditioned media. Hormones in B-G were measured on the apical side only of the transwell. In B-G, batches A and B from the RNASeq experiment were pooled so each point on the plot is the result from two organoid batches pooled together. **H**) Serotonin released from the apical or basolateral side (as indicated) from uninduced and induced *NGN3*-HIOs in response to *L. reuteri* 6475 or 17983 conditioned media. In H, shape denotes independent batches of organoids. Only p-values <0.1 are shown with p<0.05 being considered significant. Significance was determined with a Dunnett’s Test.

Interestingly, no genes related to serotonin-metabolism or transporters (*TPH1, TPH2, DDC, SLC18A1, SERT)* were altered by either *L. reuteri* strain. Nevertheless, we observed that *L. reuteri* 6475 and 17938 promote serotonin secretion (**Figure 3H**). Collectively, these data indicate that *L. reuteri* regulates numerous gut hormones; however, *L. reuteri* may upregulate either or both the expression and secretion of intestinal hormones.

### *L. reuteri* affects the transcription and secretion of enterocytic hormones

While the genes in cluster 1 were upregulated by *NGN3* induction, those in cluster 5 were downregulated by *NGN3* induction (**Figure 2**). The genes downregulated were for hormones vasopressin (AVP), adipolin (C1QTNF12), luteinizing hormone subunit B (LHB), neurotensin (NTS), and oxytocin (OXT).

Neuregulin-4 (NRG4) and tachykinin-3 (TAC3) were unaffected by induction. All these hormone genes were significantly upregulated by *L. reuteri* 6475, while only *LHB* and *OXT* were significantly upregulated by *L. reuteri* 17938. Interestingly among these hormones, only neurotensin is well established to be produced by the gut epithelium. In mice, neurotensin is observed within villus proximal enteroendocrine L-cells ^102,103^ and is thought to be produced in L cells only after they have migrated away from crypts and are exposed to increasing levels of BMP4 signaling^102^.

Recently we reported that oxytocin is produced by enterocytes in the small intestinal epithelium and its secretion is promoted by *L. reuteri*^33^. To determine if any of these hormones are also produced by enterocytes, we analyzed the adult jejunum single-cell RNA-Seq (scRNA-Seq) data within the Gut Cell Atlas^104^. While chromogranin A (*CHGA*) transcription clustered with enteroendocrine cells, transcription of *AVP*, *LHB*, and *C1QTNF12* (adipolin) clustered similarly to that for sucrose isomatase (*SI*), a marker of enterocytes (**Figure 4A-F**). Furthermore, we were able to confirm that *C1QTNF12* (adipolin) is produced in enterocytes in the human jejunum (**Figure 4G**).

**Figure 4:**
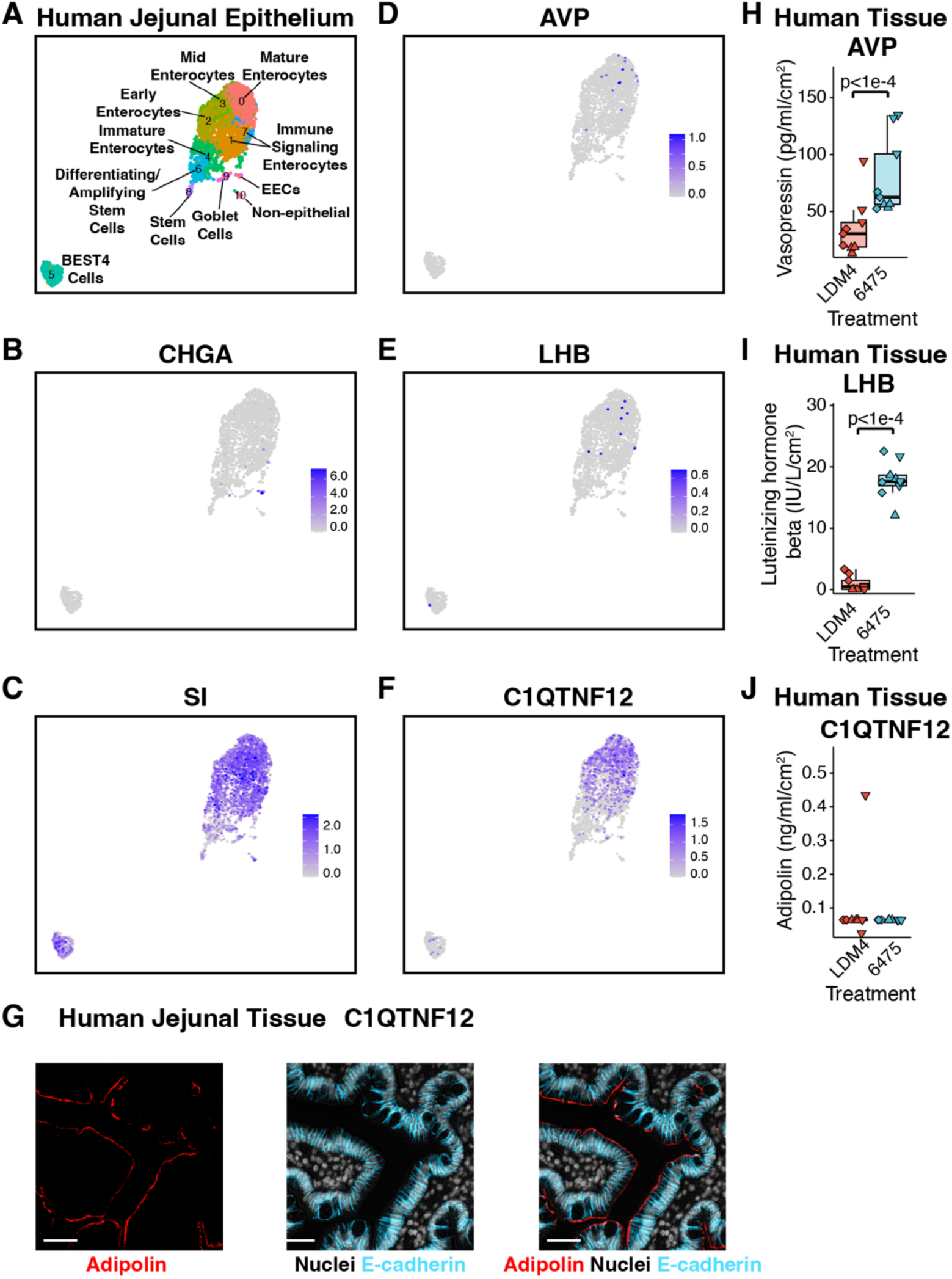
*L. reuteri* promotes the secretion of enterocytic hormones. **A)** Gut Cell Atlas annotated UMAP of the adult jejunum (adapted from Danhof et al 2023), highlighting the enteroendocrine marker *CHGA* (**B**), the enterocyte marker *SI* (**C**), vasopressin (*AVP*, **D**), luteinizing hormone subunit beta (*LHB*, **E**), and adipolin (C1QTNF12, **F**). **G**) Adipolin visualized in human jejunal tissue. Scale bar represents 50 µm. Secretion of **H**) vasopressin and **I**) luteinizing hormone subunit beta and **J**) the lack of secretion of adipolin from whole human jejunal tissue using the method shown in Figure 3A except with *ex vivo* human jejunal intestinal tissue. Shape represents unique human intestinal donors. Significance was determined using a linear mixed model with p <0.05 considered as significant.

Next we checked if *L. reuteri* is able to induce the secretion of any of these hormones from whole intestinal tissue as it does for oxytocin^33^. *L. reuteri* was able to induce the release of vasopressin and LHB but not adipolin from the human jejunum (**Figure 4H-J**). Given that *AVP* and *LHB* transcription are enriched in epithelial cells in adult gut tissue^104^ (p = 4.1e-3 for AVP in epithelium across the entire adult intestine, p = 0 for just jejunum; p = 1.0e-5 for LHB in epithelium across the entire adult intestine, p = 0.014 for just jejunum, hypergeometric distribution), the released vasopressin and LHB may originate from the epithelium rather than other regions of the intestinal tissue.

In looking at the functions of the hormones in cluster 5, these hormones have roles in sexual function and behavior, whereas those in cluster 1 have functions mostly in feeding behavior and cardiovascular function. We also noticed that kisspeptin (KISS1), a hormone characterized in the brain with roles in gonad development^105^, though not differentially regulated by *L. reuteri*, was expressed in the *NGN3*-HIOs and downregulated by induction. Like the other hormones in cluster 5, KISS1 appears to be produced in enterocytes (**Supplemental Figure 6A**). We looked to see if *L. reuteri* could induce its secretion and found no evidence of *L. reuteri* mediates release of KISS1 (**Supplemental Figure 6B**).

## Discussion

*L. reuteri* has been characterized as a beneficial microbe capable of affecting multiple aspects of host physiology within and beyond the gut. These effects are likely to involve host-microbe interactions that initiate at the intestinal epithelial layer. To begin to understand those interactions, here we used an organoid model enhanced in its number of enteroendocrine cells to specifically study interactions between *L. reuteri* and intestinal hormones. While, microbes have been identified that promote the release or expression of hormones or neuropeptides including GLP-1^106–108^, PYY^107,108^, serotonin^106,109–112^, testosterone^26^, and oxytocin^33^, our study here focused on the effect of a single microbe on intestinal hormones using a human intestinal organoid model system. Our results indicate that multiple intestinal hormones are regulated by *L. reuteri* (**Table 4**); and moreover, these data point towards there being novel hormones derived from enterocytes in the gut. Specifically, while luteinizing hormone subunit beta was previously observed in the stomach and duodenum^113^, kisspeptin, adipolin, and vasopressin have not been described as intestinal epithelial hormones.

**Table 4:** Summary of *L. reuteri’s* effects on gut hormones. +, upregulated; -, downregulated; ND, not determined; NS, not significant; *not confirmed if secretion occurs from epithelial cells

| Hormone | Proposed or established hormone cell type | Expression | Secretion |
| --- | --- | --- | --- |
| Amylin | Enteroendocrine | ND | + (this work) |
| C-peptide | Enteroendocrine | ND | + (this work) |
| CCK | Enteroendocrine | + (this work) | ND |
| Gastrin | Enteroendocrine | + (this work) | ND |
| Ghrelin | Enteroendocrine | + (this work) | NS (this work) |
| GIP | Enteroendocrine | + (this work) | + (this work) |
| Luteinizing hormone, beta subunit | Enterocyte | + (this work) | + (this work)* |
| Motilin | Enteroendocrine | + (this work) | ND |
| Neurotensin | Enteroendocrine | + (this work) | ND |
| NPW | Enteroendocrine | + (this work) | ND |
| NPY | Enteroendocrine | + (this work) | ND |
| Oxytocin | Enterocyte | + (this work) | +<sup>33</sup> |
| PYY | Enteroendocrine | NS (this work) | + (this work) |
| Secretin | Enteroendocrine | NS (this work) | +<sup>33</sup> |
| Serotonin | Enteroendocrine | NS (this work) | + (this work) |
| Somatostatin | Enteroendocrine | + (this work) | ND |
| Vasopressin | Enterocyte | + (this work) | + (this work)* |

While we found several well-known intestinal hormones are not regulated by *L. reuteri* (including GLP-1 and pancreatic peptide (PP), we observed that *L. reuteri* largely transcriptionally upregulates gut hormones. We also found that a smaller set of gut hormones is secreted by *L. reuteri*. This study was particularly focused on the effect of *L. reuteri* on hormones of the small intestine, where we postulate *L. reuteri* may act therapeutically in humans. Hence, these data broadly suggest that *L. reuteri* can potentially act beneficially via regulation of intestinal hormones. Moreover, our study considered not just a single probiotic strain of *L. reuteri* but two different commercially used strains. Interestingly, our study failed to observe major differences between the two strains: *L. reuteri* 17938 appeared to transcriptionally affect HIOs enriched in enteroendocrine cells very similarly to *L. reuteri* 6475, albeit with a lower magnitude. Furthermore, the select hormones whose secretion we tested were similarly induced by both strains. An unknown experimental condition could be responsible for *L. reuteri* 17938’s lower effect on the HIO transcripts.

Recently, several new enteric hormones have been described. In addition to the discovery of oxytocin in the intestinal epithelium, famsin^114^, GDF15^115^, and cholesin^116^ have been discovered. A survey of these peptide hormones in the Gut Cell Atlas^104^ suggests that, in addition to the previously described FGF19, guanylin, and uroguanylin^31^, these hormones are made in enterocytes rather than enteroendocrine cells. The recognition that enterocytes can produce hormones has opened questions regarding the production of these hormones. Enteroendocrine cell-derived hormones are produced from prohormones that are cleaved to the active hormone by prohormone convertases some of which are exclusively produced in enteroendocrine cells^117^ and are subsequently secreted from vesicles stored in axon-like structures within the cell^118^ on stimulation. Hence, are these enterocytic hormones only processed by convertases that are made in enterocytes? Are the hormones stored in vesicles like in enteroendocrine cells? And how and to where are these vesicles released?

The function of these novel enterocytic hormones is additionally waiting to be determined. Interestingly, non-intestinal sources of oxytocin, vasopressin, kisspeptin, and luteinizing hormone have roles in regulating sexual function, and several also function in regulating eating or digestion. Famsin^114^, GDF15^115^, and cholesin^116^ have been characterized with roles related to metabolism and energy regulation. Given the known links between metabolic state and sexual function^119^, potentially then, intestinal sources of oxytocin, vasopressin, kisspeptin, and luteinizing hormone serve to link metabolic state to sexual function.

We also observed that adipolin is produced in the small intestinal epithelial layer. Adipolin has been observed as present in the small intestinal epithelium presented by the Human Protein Atlas (https://www.proteinatlas.org/ENSG00000184163-C1QTNF12/tissue/small+intestine)^120^. In adipose tissue, adipolin was characterized as an adipokine that improves glucose tolerance and insulin response and reduces macrophages and proinflammatory immune responses^86^. In the intestine, it may have similar immune and metabolic functions.

Previously we determined that the hormone secretin is involved in *L. reuteri’s* release of oxytocin^33^. However, what *L. reuteri* makes to promote secretin’s release is currently unknown. Presently, a variety of different microbial metabolites or structures have been shown to promote the release of or are associated with the release of intestinal hormones. These include short chain fatty acids^121–123^, branched and aromatic amino acids^123^, indoles^124^, secondary bile acids^125^, and microvesicles^112^. Whether any of these molecules or others produced by *L. reuteri* are involved in the hormones affected here remains to be determined.

A few limitations of our study design should be mentioned. First, the media conditions of the organoids have been observed to reduce inflammatory responses^126^. Second, the organoids only represent the epithelial layer so interactions between *L. reuteri* and the host that depend on immune cells, enteric neurons, or products of the lamina propria or circulation cannot be captured by this assay. Third, the assay was performed using cell-free supernatants with a three-hour exposure. Hence, host responses that require intact structural components of *L. reuteri* or a different length of exposure are also not represented in this assay. Fourth, the secretion assays were not designed to capture whether *L. reuteri* suppresses the secretion of hormones, and similarly the transcriptomic data only considers *L.* reuteri’s effect relative to bacterial growth media. Further follow-up studies will be needed to determine if *L. reuteri* is able to promote secretion of these hormones under more physiologically relevant conditions.

In conclusion, this work demonstrates that *L. reuteri* regulates several canonical and novel hormones of the intestinal epithelial layer. These results open exciting investigations regarding how *L. reuteri* may influence a wide range of aspects of systemic physiology.

## Acknowledgements

We would like to thank Susan Venable, Colleen Ardis, Javier Nieto, and the LifeGift donor families for their assistance and support during this project. This project was supported in part by NIH grant DK056338 (Cellular and Molecular Core, Functional Genomics and Microbiome Core, and Gastrointestinal Experimental Model Systems Core), which supports the Texas Medical Center Digestive Diseases Center. This project was supported in part by the Optical Imaging and Vital Microscopy Core at Baylor College of Medicine.

---

**Supplemental Figure S1.**
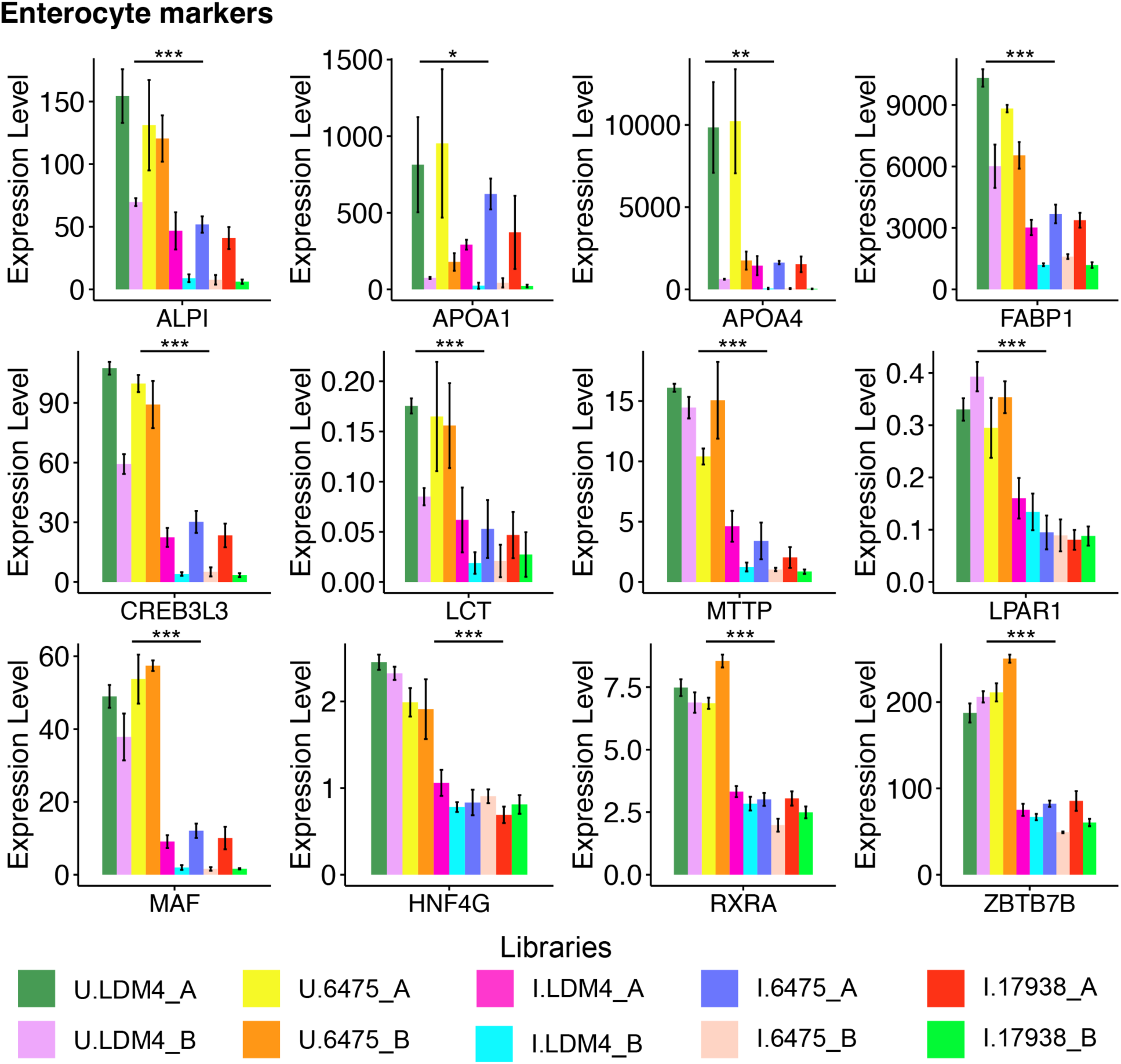
Expression levels of enterocyte cell markers (*ALPI*, *APOA1*, *APOA4*, *FABP1*, *CREB3L3*, *LCT*, *MTTP*, *LPAR1*, *MAF*, *HNF4G*, *RXRA*, *ZBTB7B*) in uninduced and induced NGN3-HIOs. Libraries are labeled “U” for uninduced, “I” for induced” and “A” and “B” for the first and second batches of *NGN3*-HIOs. Expression levels shown are counts per million GeTMM transformed read counts. Significance of expression levels between the uninduced and induced libraries was calculated using a two-sample, two-sided, Mann-Whitney test. *, p<0.05, **, p<0.005, ***, p<0.0005.

**Supplemental Figure S2.**
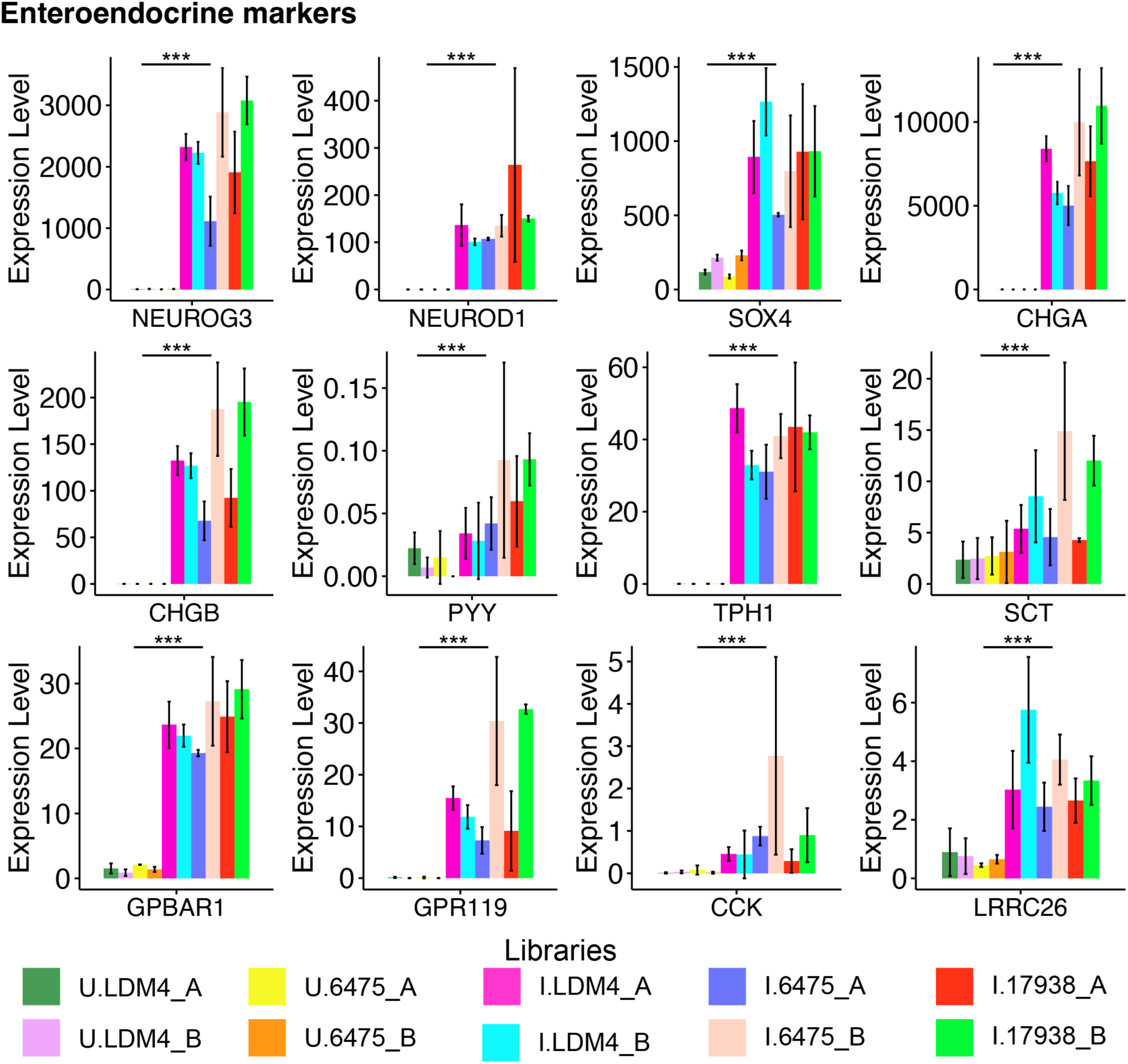
Expression levels of enteroendocrine cell precursor markers (*NEUROG3*, *NEUROD1*, *SOX4*) and cell markers (*CHGA*, *CHGB*, *PYY*, *TPH1*, *SCT*, *GPBAR1*, *GPT119*, *CCK*, *LRRC26*) in uninduced and induced *NGN3*-HIOs. Libraries are labeled “U” for uninduced, “I” for induced” and “A” and “B” for the first and second batches of *NGN3*-HIOs. Expression levels shown are counts per million GeTMM transformed read counts. Significance of expression levels between the uninduced and induced libraries was calculated using a two-sample, two-sided, Mann-Whitney test. *, p<0.05, **, p<0.005, ***, p<0.0005.

**Supplemental Figure S3.**
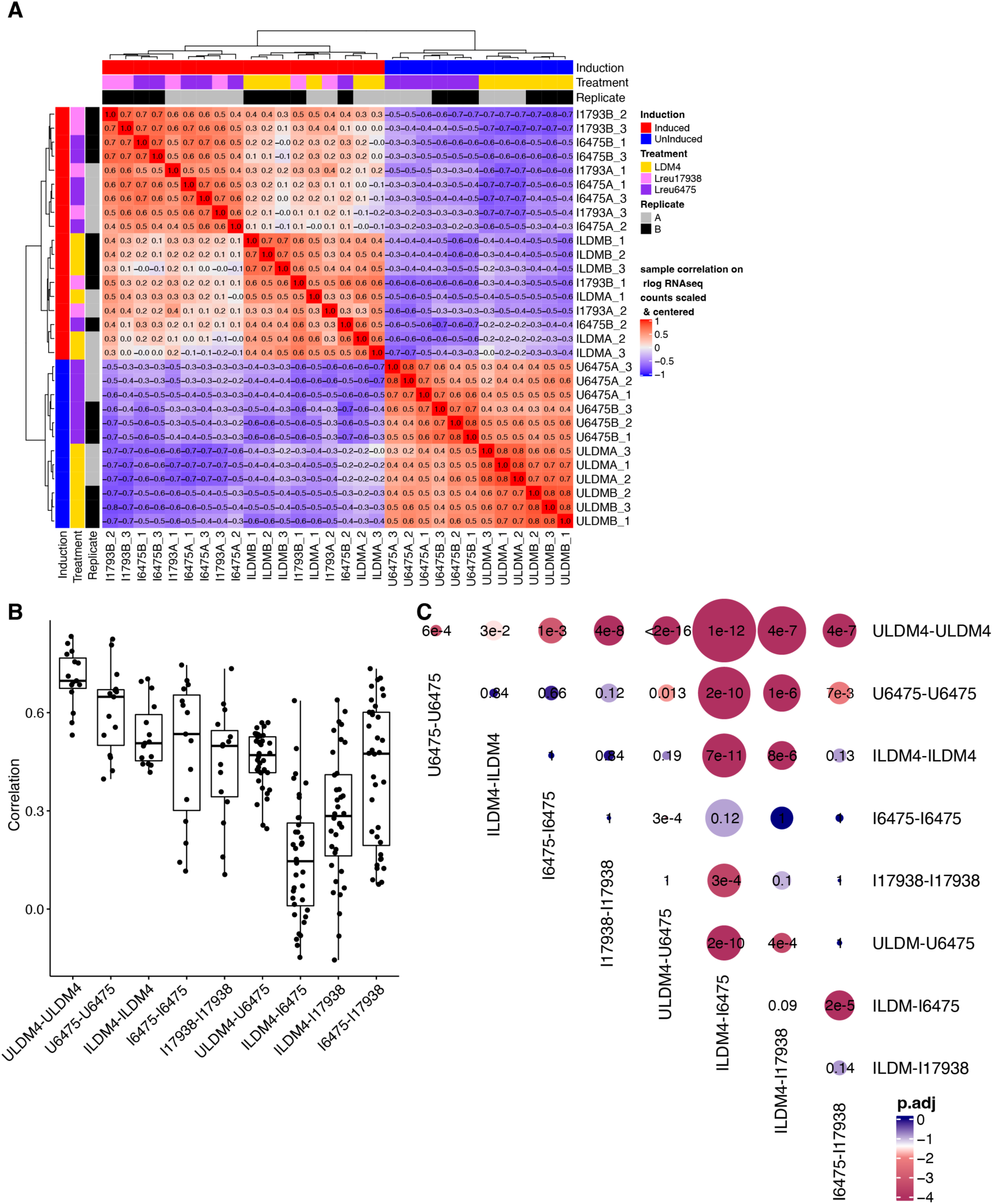
Similarity and differences among *NGN3*-HIO transcriptomes. **A)** Boxplots of Pearson correlation values within and between transcriptomes. **B)** Correlogram of mean differences (circle size) and adjusted p-values (circle fill) between comparisons shown in A. Samples are listed, whereby “U” refers to uninduced *NGN3*-HIOs, “I” for induced *NGN3*-HIOs, “LDM4” for media only treatment, “6475” for *L. reuteri* 6475 treatment, “17938” for *L. reuteri* 17938 treatment, “A” or “B” refers to the biological replicate, and “1”, “2”, or “3” refers to the technical replicate within each biological replicate. **C)** Genes differentially regulated between *L. reuteri* 6475 and 17938 on induced *NGN3*-HIOs. The graph shows the log_2_ fold change expression of the gene for the indicated comparison. The bars are colored using the log_10_ scaled mean GeTMM counts to illustrate how abundantly expressed the gene is. Transparent overlays are used on genes not differentially expressed for the given comparison. Comparisons shown: U6475-ULDM4, *L. reuteri* 6475 on uninduced HIOs compared to LDM4 media control; I6475-ILDM4, *L. reuteri* 6475 on induced HIOs compared to LDM4 media control; I17938- ILDM4, *L. reuteri* 17938 on induced HIOs compared to LDM4 media control; I6475-I17938 *L. reuteri* 6475 compared to *L. reuteri* 17938 on induced HIOs; ILDM4-ULDM4, LDM4 media control on induced versus uninduced HIOs; I6475-U6475, *L. reuteri* 6475 on induced versus uninduced HIOs. For each, positive fold changes indicate genes upregulated by the condition listed first.

**Supplemental Figure S4.**
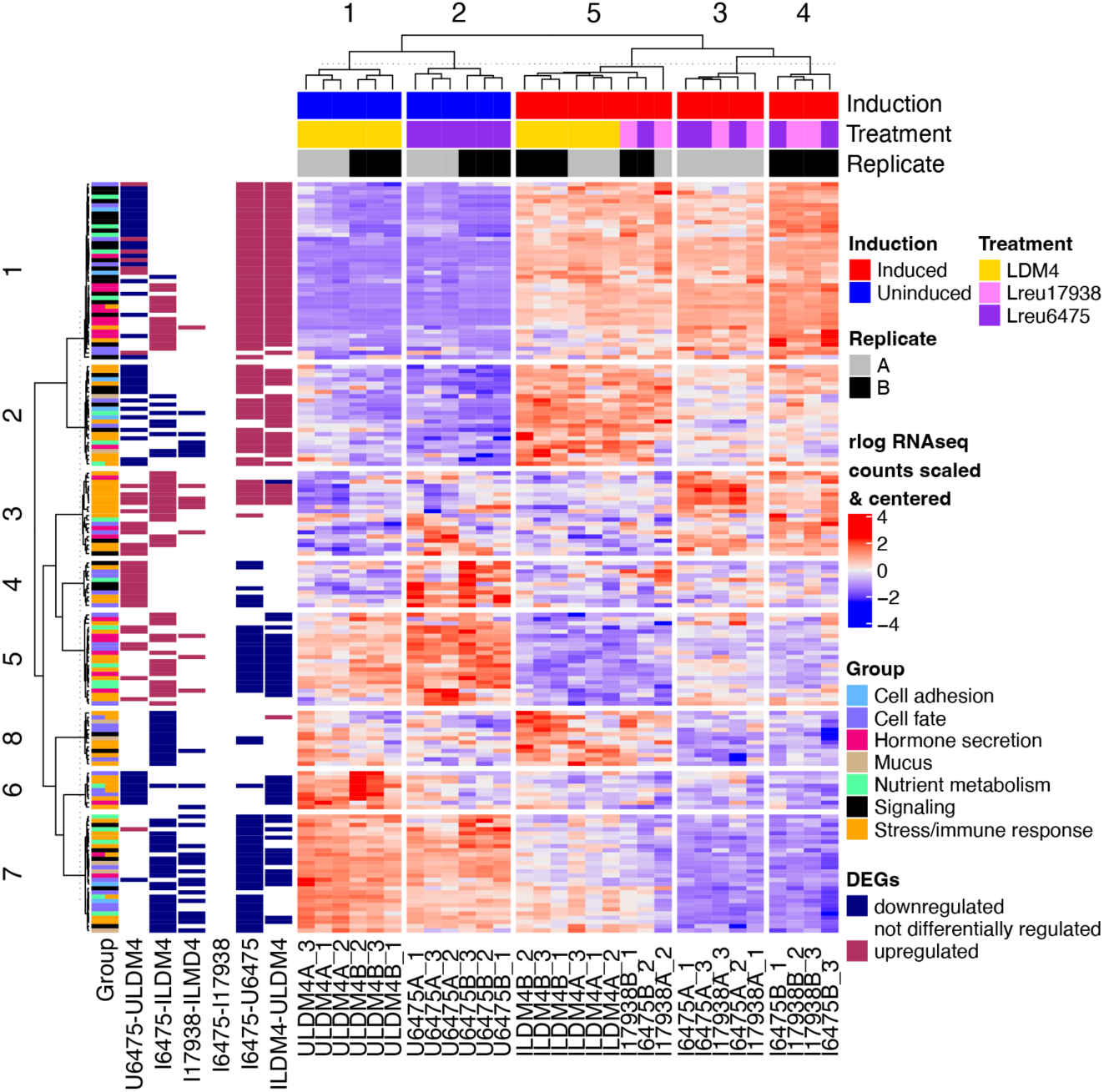
Cluster analysis of DEGs belonging to functionally enriched groups. The heatmap shows gene expression values as rlog counts that were scaled and centered. Samples (the columns) along the bottom of the heatmap are labeled as “U” for uninduced *NGN3*-HIOs, “I” for induced *NGN3*-HIOs, “LDM4” for media only treatment, “6475” for *L. reuteri* 6475 treatment, “17938” for *L. reuteri* 17938 treatment, “A” or “B” for the biological replicate, and “1”, “2”, or “3” for the technical replicate within each biological replicate. Samples are annotated above the heatmap as shown in the legend. Genes (rows) were arranged by K-means clustering and annotated into groups as shown in the legend. For each sample comparison (e.g. U6475-ULDM4), if the gene was down or upregulated (e.g. higher in U6475 than ULDM4), a color is given as shown in the legend.

**Supplemental Figure S5.**
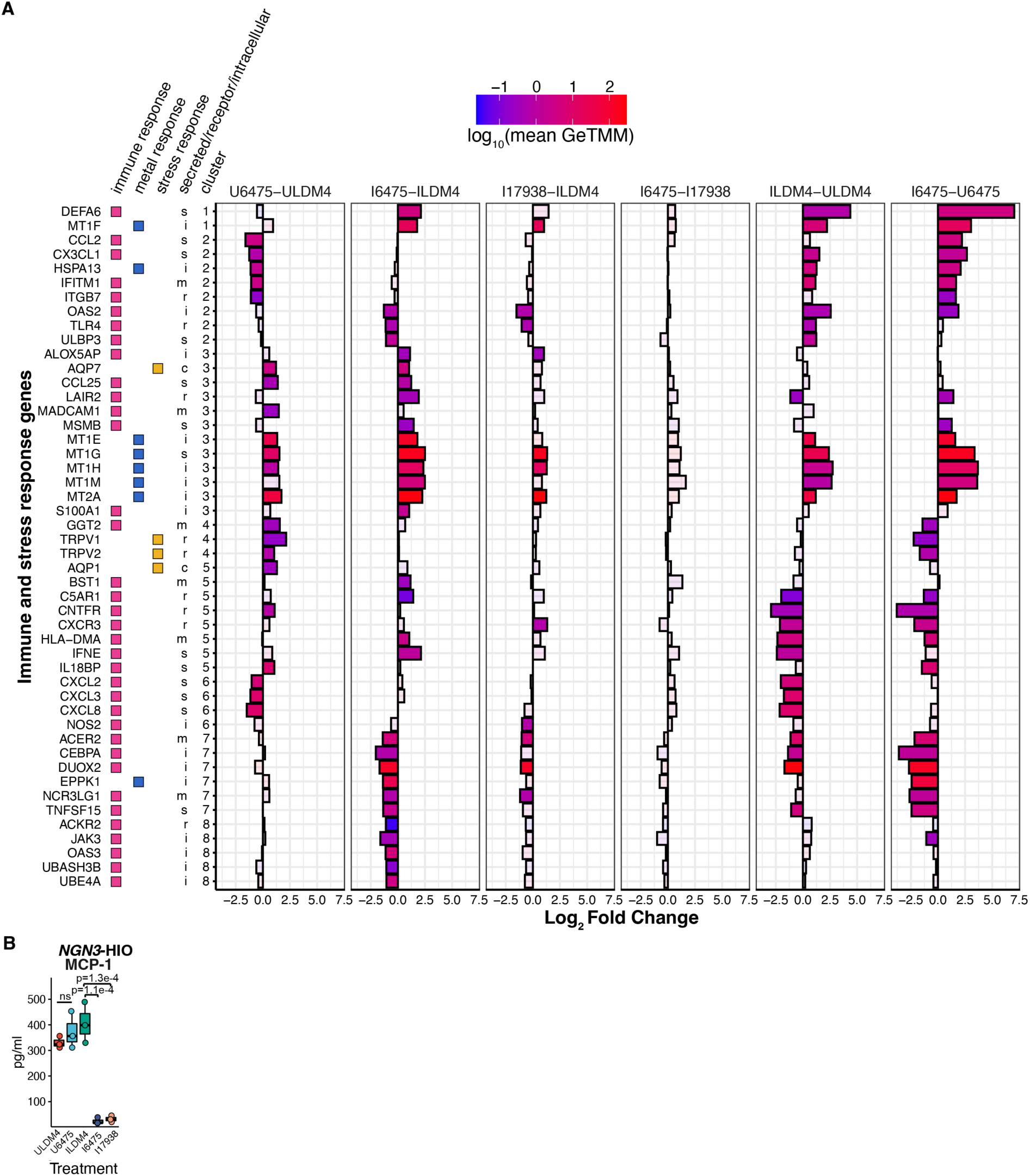
*L. reuteri* regulates immune, metal, and stress response. **A)** Immune, metal, and stress genes differentially regulated by *L. reuteri*. The genes are annotated with their function, whether they are secreted, a receptor, or intercellular, and what cluster they belong to relative to Supplemental Figure S4. The graph shows the log_2_ fold change expression of the gene for the indicated comparison. The bars are colored using the log_10_ scaled mean GeTMM counts to illustrate how abundantly expressed the gene is. Transparent overlays are used on genes not differentially expressed for the given comparison. Comparisons shown: U6475-ULDM4, *L. reuteri* 6475 on uninduced HIOs compared to LDM4 media control; I6475-ILDM4, *L. reuteri* 6475 on induced HIOs compared to LDM4 media control; I17938-ILDM4, *L. reuteri* 17938 on induced HIOs compared to LDM4 media control; I6475-I17938 *L. reuteri* 6475 compared to *L. reuteri* 17938 on induced HIOs; ILDM4-ULDM4, LDM4 media control on induced versus uninduced HIOs; I6475-U6475, *L. reuteri* 6475 on induced versus uninduced HIOs. For each, positive fold changes indicate genes upregulated by the condition listed first. **B)** MCP-1 protein levels measured by Luminex on uninduced (U) or induced (I) HIOs treated with *L. reuteri* 6475 or 17938.

**Supplemental Figure S6:**
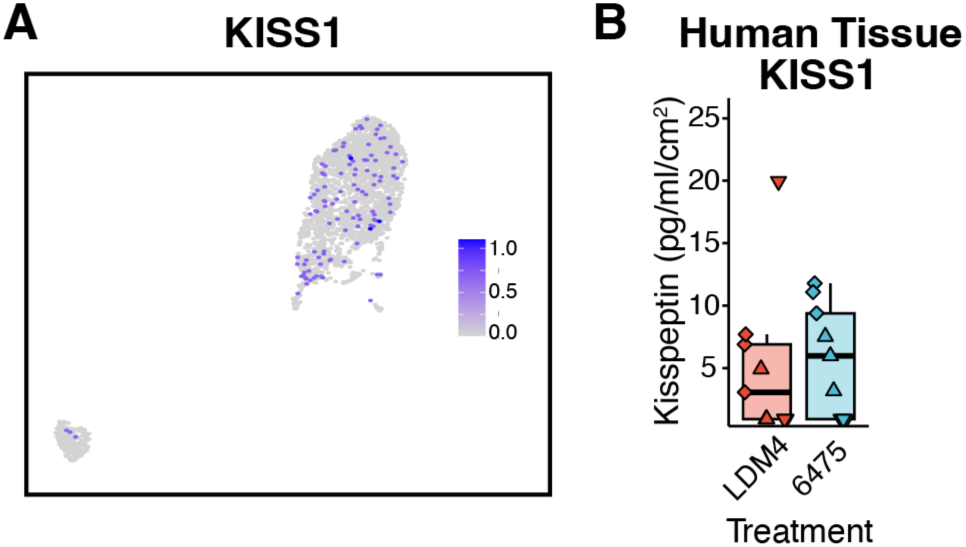
KISS1 may be produced in the intestinal epithelium. **A)** UMAP of KISS1 using the Gut Cell Atlas adult jejunum data. **B)** Lack of secretion of kisspeptin in response to bacterial media control (LDM4) and *L. reuteri* 6475 conditioned media from *ex vivo* human jejunal intestinal tissue. Shape represents unique human intestinal donors. Significance was determined using a linear mixed model with p<0.05 considered as significant.

## References

1. Savino, F., Pelle, E., Palumeri, E., Oggero, R. & Miniero, R. Lactobacillus reuteri (American Type Culture Collection Strain 55730) Versus Simethicone in the Treatment of Infantile Colic: A Prospective Randomized Study. Pediatrics 119, e124–e130 (2007).

2. Nilsson, A. G., Sundh, D., Bäckhed, F. & Lorentzon, M. Lactobacillus reuteri reduces bone loss in older women with low bone mineral density: a randomized, placebo-controlled, double-blind, clinical trial. J. Intern. Med. 284, 307–317 (2018).

3. Thomas, C. M. et al. Histamine derived from probiotic Lactobacillus reuteri suppresses TNF via modulation of PKA and ERK signaling. PloS One 7, e31951 (2012).

4. Walter, J., Britton, R. A. & Roos, S. Host-microbial symbiosis in the vertebrate gastrointestinal tract and the Lactobacillus reuteri paradigm. Proc. Natl. Acad. Sci. U. S. A. 108 **Suppl 1**, 4645–4652 (2011).

5. Lin, Y. P., Thibodeaux, C. H., Peña, J. A., Ferry, G. D. & Versalovic, J. Probiotic Lactobacillus reuteri suppress proinflammatory cytokines via c-Jun. Inflamm. Bowel Dis. 14, 1068–1083 (2008).

6. Jones, S. E. & Versalovic, J. Probiotic Lactobacillus reuteri biofilms produce antimicrobial and anti-inflammatory factors. BMC Microbiol. 9, 35 (2009).

7. Buffington, S. A. et al. Microbial Reconstitution Reverses Maternal Diet- Induced Social and Synaptic Deficits in Offspring. Cell 165, 1762–1775 (2016).

8. Sgritta, M. et al. Mechanisms Underlying Microbial-Mediated Changes in Social Behavior in Mouse Models of Autism Spectrum Disorder. Neuron 101, 246–259.e6 (2019).

9. Buffington, S. A. et al. Dissecting the contribution of host genetics and the microbiome in complex behaviors. Cell 184, 1740–1756.e16 (2021).

10. Schmitt, L. M. et al. Results of a phase Ib study of SB-121, an investigational probiotic formulation, a randomized controlled trial in participants with autism spectrum disorder. Sci. Rep. 13, 5192 (2023).

11. Mazzone, L. et al. Precision microbial intervention improves social behavior but not autism severity: A pilot double-blind randomized placebo-controlled trial. Cell Host Microbe 32, 106–116.e6 (2024).

12. Rosander, A., Connolly, E. & Roos, S. Removal of Antibiotic Resistance Gene-Carrying Plasmids from Lactobacillus reuteri ATCC 55730 and Characterization of the Resulting Daughter Strain, L. reuteri DSM 17938. Appl. Environ. Microbiol. 74, 6032–6040 (2008).

13. Spinler, J. K. et al. From Prediction to Function Using Evolutionary Genomics: Human- Specific Ecotypes of Lactobacillus reuteri Have Diverse Probiotic Functions. Genome Biol. Evol. 6, 1772–1789 (2014).

14. Indrio, F. et al. The Effects of Probiotics on Feeding Tolerance, Bowel Habits, and Gastrointestinal Motility in Preterm Newborns. J. Pediatr. 152, 801–806 (2008).

15. Coccorullo, P. et al. Lactobacillus reuteri (DSM 17938) in Infants with Functional Chronic Constipation: A Double-Blind, Randomized, Placebo-Controlled Study. J. Pediatr. 157, 598–602 (2010).

16. Miniello, V. L. et al. Lactobacillus reuteri modulates cytokines production in exhaled breath condensate of children with atopic dermatitis. J. Pediatr. Gastroenterol. Nutr. 50, 573–576 (2010).

17. Quach, D., Parameswaran, N., McCabe, L. & Britton, R. A. Characterizing how probiotic Lactobacillus reuteri 6475 and lactobacillic acid mediate suppression of osteoclast differentiation. Bone Rep. 11, 100227 (2019).

18. Rios-Arce, N. D. et al. Post-antibiotic gut dysbiosis-induced trabecular bone loss is dependent on lymphocytes. Bone 134, 115269 (2020).

19. Schepper, J. D. et al. Involvement of the Gut Microbiota and Barrier Function in Glucocorticoid-Induced Osteoporosis. J. Bone Miner. Res. Off. J. Am. Soc. Bone Miner. Res. 35, 801–820 (2020).

20. Schepper, J. D. et al. Probiotic Lactobacillus reuteri Prevents Postantibiotic Bone Loss by Reducing Intestinal Dysbiosis and Preventing Barrier Disruption. J. Bone Miner. Res. Off. J. Am. Soc. Bone Miner. Res. 34, 681–698 (2019).

21. Zhang, J. et al. Loss of Bone and Wnt10b Expression in Male Type 1 Diabetic Mice Is Blocked by the Probiotic Lactobacillus reuteri. Endocrinology 156, 3169–3182 (2015).

22. Britton, R. A. et al. Probiotic L. reuteri treatment prevents bone loss in a menopausal ovariectomized mouse model. J. Cell. Physiol. 229, 1822–1830 (2014).

23. McCabe, L. R., Irwin, R., Schaefer, L. & Britton, R. A. Probiotic use decreases intestinal inflammation and increases bone density in healthy male but not female mice: *L. reuteri* PROMOTES INTESTINE AND BONE HEALTH. J. Cell. Physiol. 228, 1793–1798 (2013).

24. Poutahidis, T. et al. Microbial Symbionts Accelerate Wound Healing via the Neuropeptide Hormone Oxytocin. PLOS ONE 8, e78898 (2013).

25. Erdman, S. & Poutahidis, T. Probiotic ‘glow of health’: it’s more than skin deep. Benef. Microbes 5, 109–119 (2014).

26. Poutahidis, T. et al. Probiotic Microbes Sustain Youthful Serum Testosterone Levels and Testicular Size in Aging Mice. PLoS ONE 9, e84877 (2014).

27. Mu, Q., Tavella, V. J. & Luo, X. M. Role of Lactobacillus reuteri in Human Health and Diseases. Front. Microbiol. 9, (2018).

28. Saulnier, D. M. et al. Exploring Metabolic Pathway Reconstruction and Genome-Wide Expression Profiling in Lactobacillus reuteri to Define Functional Probiotic Features. PLoS ONE 6, (2011).

29. Liu, Y. et al. Probiotic-Derived Ecto-5’-Nucleotidase Produces Anti-Inflammatory Adenosine Metabolites in Treg-Deficient Scurfy Mice. Probiotics Antimicrob. Proteins 15, 1001–1013 (2023).

30. Worthington, J. J., Reimann, F. & Gribble, F. M. Enteroendocrine cells-sensory sentinels of the intestinal environment and orchestrators of mucosal immunity. Mucosal Immunol. 11, 3– 20 (2018).

31. Bany Bakar, R., Reimann, F. & Gribble, F. M. The intestine as an endocrine organ and the role of gut hormones in metabolic regulation. Nat. Rev. Gastroenterol. Hepatol. 20, 784–796 (2023).

32. Chang-Graham, A. L. et al. Human Intestinal Enteroids With Inducible Neurogenin-3 Expression as a Novel Model of Gut Hormone Secretion. Cell. Mol. Gastroenterol. Hepatol. 8, 209–229 (2019).

33. Danhof, H. A., Lee, J., Thapa, A., Britton, R. A. & Di Rienzi, S. C. Microbial stimulation of oxytocin release from the intestinal epithelium via secretin signaling. Gut Microbes 15, 2256043 (2023).

34. Illumina Inc. Sequencing Analysis Software User Guide for Pipeline Version 1.3 and CASAVA Version 1.0 Illumina Inc. (San Diego, CA, USA, 2008).

35. Dobin, A. et al. STAR: ultrafast universal RNA-seq aligner. Bioinformatics 29, 15–21 (2013).

36. Anders, S., Pyl, P. T. & Huber, W. HTSeq—a Python framework to work with high- throughput sequencing data. Bioinformatics 31, 166–169 (2015).

37. Ge, S. X., Son, E. W. & Yao, R. iDEP: an integrated web application for differential expression and pathway analysis of RNA-Seq data. BMC Bioinformatics 19, 534 (2018).

38. Oksanen, J. et al. vegan: Community Ecology Package. (2019).

39. Gu, Z., Eils, R. & Schlesner, M. Complex heatmaps reveal patterns and correlations in multidimensional genomic data. Bioinformatics (2016).

40. Kassambara, A. Ggpubr: ‘ggplot2’ Based Publication Ready Plots. (2019).

41. Holm, S. A Simple Sequentially Rejective Multiple Test Procedure. Scand. J. Stat. 6, 65–70 (1979).

42. Love, M. I., Huber, W. & Anders, S. Moderated estimation of fold change and dispersion for RNA-seq data with DESeq2. Genome Biol. 15, 550 (2014).

43. Benjamini, Y. & Hochberg, Y. Controlling the False Discovery Rate: A Practical and Powerful Approach to Multiple Testing. J. R. Stat. Soc. Ser. B Methodol. 57, 289–300 (1995).

44. Zerbino, D. R. et al. Ensembl 2018. Nucleic Acids Res. 46, D754–D761 (2018).

45. Fabregat, A. et al. The Reactome Pathway Knowledgebase. Nucleic Acids Res. 46, D649– D655 (2018).

46. Jassal, B. et al. The reactome pathway knowledgebase. Nucleic Acids Res. gkz1031, 1–6 (2019).

47. Mi, H. et al. Protocol Update for large-scale genome and gene function analysis with the PANTHER classification system (v.14.0). Nat. Protoc. 14, 703–721 (2019).

48. Stelzer, G. et al. The GeneCards Suite: From Gene Data Mining to Disease Genome Sequence Analyses. Curr. Protoc. Bioinforma. 54, (2016).

49. Ritchie, M. E. et al. limma powers differential expression analyses for RNA-sequencing and microarray studies. Nucleic Acids Res. 43, e47–e47 (2015).

50. R Core Team. R: A Language and Environment for Statistical Computing. (R Foundation for Statistical Computing, Vienna, Austria, 2018).

51. Smid, M. et al. Gene length corrected trimmed mean of M-values (GeTMM) processing of RNA-seq data performs similarly in intersample analyses while improving intrasample comparisons. BMC Bioinformatics 19, 236 (2018).

52. Robinson, M. D., McCarthy, D. J. & Smyth, G. K. edgeR: a Bioconductor package for differential expression analysis of digital gene expression data. Bioinforma. Oxf. Engl. 26, 139–140 (2010).

53. Haber, A. L. et al. A single-cell survey of the small intestinal epithelium. Nature 551, 333– 339 (2017).

54. Reaux, A., Fournie-Zaluski, M. C. & Llorens-Cortes, C. Angiotensin III: a central regulator of vasopressin release and blood pressure. Trends Endocrinol. Metab. 12, 157–162 (2001).

55. Szczepańska-Sadowska, E. Interaction of vasopressin and angiotensin II in central control of blood pressure and thirst. Regul. Pept. 66, 65–71 (1996).

56. van Unen, J. et al. Kinetics of recruitment and allosteric activation of ARHGEF25 isoforms by the heterotrimeric G-protein Gαq. Sci. Rep. 6, 36825 (2016).

57. Coate, K. C., Kliewer, S. A. & Mangelsdorf, D. J. SnapShot: Hormones of the Gastrointestinal Tract. Cell 159, 1478–1478.e1 (2014).

58. Luyer, M. D. et al. Nutritional stimulation of cholecystokinin receptors inhibits inflammation via the vagus nerve. J. Exp. Med. 202, 1023–1029 (2005).

59. Lovick, T. A. CCK as a modulator of cardiovascular function. J. Chem. Neuroanat. 38, 176– 184 (2009).

60. Mace, O. J., Tehan, B. & Marshall, F. Pharmacology and physiology of gastrointestinal enteroendocrine cells. Pharmacol. Res. Perspect. 3, e00155 (2015).

61. Gutzwiller, J.-P. et al. Effect of intravenous human gastrin-releasing peptide on food intake in humans. Gastroenterology 106, 1168–1173 (1994).

62. Russo, F. et al. The obestatin/ghrelin ratio and ghrelin genetics in adult celiac patients before and after a gluten-free diet, in irritable bowel syndrome patients and healthy individuals: *Eur*. J. Gastroenterol. Hepatol. 29, 160–168 (2017).

63. Malik, S., McGlone, F., Bedrossian, D. & Dagher, A. Ghrelin Modulates Brain Activity in Areas that Control Appetitive Behavior. Cell Metab. 7, 400–409 (2008).

64. Zhang, G. et al. Ghrelin and Cardiovascular Diseases. Curr. Cardiol. Rev. 6, 62–70 (2010).

65. Li, H. et al. Gastric neuropeptide W is regulated by meal-related nutrients. Peptides 62, 6–14 (2014).

66. Baker, J. R., Cardinal, K., Bober, C., Taylor, M. M. & Samson, W. K. Neuropeptide W acts in brain to control prolactin, corticosterone, and growth hormone release. Endocrinology 144, 2816–2821 (2003).

67. Levine, A. S., Winsky-Sommerer, R., Huitron-Resendiz, S., Grace, M. K. & de Lecea, L. Injection of neuropeptide W into paraventricular nucleus of hypothalamus increases food intake. Am. J. Physiol. Regul. Integr. Comp. Physiol. 288, R1727–1732 (2005).

68. Manfredi-Lozano, M., Roa, J. & Tena-Sempere, M. Connecting metabolism and gonadal function: Novel central neuropeptide pathways involved in the metabolic control of puberty and fertility. Front. Neuroendocrinol. 48, 37–49 (2018).

69. Lim, Ramon. Neuropeptide Y in Brain Function. in Handbook of Neurochemistry and Molecular Neurobiology: Neuroactive Proteins and Peptides 524–543 (Springer US, New York, NY, 2006).

70. Gribble, F. M. & Reimann, F. Enteroendocrine Cells: Chemosensors in the Intestinal Epithelium. Annu. Rev. Physiol. 78, 277–299 (2016).

71. Carroll, R. Endocrine System. in Elsevier’s Integrated Physiology 157–176 (Elsevier Health Sciences, 2007).

72. Saras, J., Grönberg, M., Stridsberg, M., Oberg, K. E. & Janson, E. T. Somatostatin induces rapid contraction of neuroendocrine cells. FEBS Lett. 581, 1957–1962 (2007).

73. Serio, R. & Zizzo, M. G. The multiple roles of dopamine receptor activation in the modulation of gastrointestinal motility and mucosal function. Auton. Neurosci. 244, 103041 (2023).

74. Villarroya, F., Cereijo, R., Villarroya, J. & Giralt, M. Brown adipose tissue as a secretory organ. Nat. Rev. Endocrinol. 13, 26–35 (2017).

75. Wang, Y., Huang, S. & Yu, P. Association between circulating neuregulin4 levels and diabetes mellitus: A meta-analysis of observational studies. PLOS ONE 14, e0225705 (2019).

76. Temur, M. et al. Increased serum neuregulin 4 levels in women with polycystic ovary syndrome: A case-control study. Ginekol. Pol. 88, 517–522 (2017).

77. Akema, T., Praputpittaya, C. & Kimura, F. Effects of Preoptic Microinjection of Neurotensin on Luteinizing Hormone Secretion in Unanesthetized Ovariectomized Rats with or without Estrogen Priming. Neuroendocrinology 46, 345–349 (1987).

78. Prague, J. K. & Dhillo, W. S. Neurokinin 3 receptor antagonism – the magic bullet for hot flushes? Climacteric 20, 505–509 (2017).

79. Lim, Ramon. Handbook of Neurochemistry and Molecular Neurobiology: Neuroactive Proteins and Peptides. (Springer US, New York, NY, 2006).

80. Nielsen, S. et al. Aquaporins in the Kidney: From Molecules to Medicine. Physiol. Rev. 82, 205–244 (2002).

81. Robertson, G. L. Abnormalities of thirst regulation. Kidney Int. 25, 460–469 (1984).

82. Thornton, S. N. Thirst and hydration: Physiology and consequences of dysfunction. Physiol. Behav. 100, 15–21 (2010).

83. Carroll, H. A. & James, L. J. Hydration, Arginine Vasopressin, and Glucoregulatory Health in Humans: A Critical Perspective. Nutrients 11, 1201 (2019).

84. Lim, Ramon. Oxytocin and Vasopressin: Genetics and Behavioral Implications. in *Handbook of Neurochemistry and Molecular Neurobiology: Neuroactive Proteins and Peptides* 574– 607 (Springer US, New York, NY, 2006).

85. Walum, H. & Young, L. J. The neural mechanisms and circuitry of the pair bond. Nat. Rev. Neurosci. 19, 643–654 (2018).

86. Enomoto, T. et al. Adipolin/C1qdc2/CTRP12 protein functions as an adipokine that improves glucose metabolism. J. Biol. Chem. 286, 34552–34558 (2011).

87. Barbe, A., et al. Adipolin (C1QTNF12) is a new adipokine in female reproduction: expression and function in porcine granulosa cells. Reprod. Camb. Engl. 167, e230272 (2024).

88. Lee, S. L. et al. Luteinizing hormone deficiency and female infertility in mice lacking the transcription factor NGFI-A (Egr-1). Science 273, 1219–1221 (1996).

89. Lofrano-Porto Adriana et al. Luteinizing Hormone Beta Mutation and Hypogonadism in Men and Women. N. Engl. J. Med. 357, 897–904 (2007).

90. Jacob, S. et al. Association of the oxytocin receptor gene (OXTR) in Caucasian children and adolescents with autism. Neurosci. Lett. 417, 6–9 (2007).

91. Welch, M. G., Margolis, K. G., Li, Z. & Gershon, M. D. Oxytocin regulates gastrointestinal motility, inflammation, macromolecular permeability, and mucosal maintenance in mice. Am. J. Physiol. - Gastrointest. Liver Physiol. 307, G848–G862 (2014).

92. Afroze, S. et al. The physiological roles of secretin and its receptor. Ann. Transl. Med. 1, 29 (2013).

93. Zhu, Y., Bond, J. & Thomas, P. Identification, classification, and partial characterization of genes in humans and other vertebrates homologous to a fish membrane progestin receptor. Proc. Natl. Acad. Sci. U. S. A. 100, 2237–2242 (2003).

94. Hwang, S. J. et al. P2Y1 purinoreceptors are fundamental to inhibitory motor control of murine colonic excitability and transit. J. Physiol. 590, 1957–1972 (2012).

95. Ma, J. et al. Glycogen metabolism regulates macrophage-mediated acute inflammatory responses. Nat. Commun. 11, 1769 (2020).

96. Bates, D., Mächler, M., Bolker, B. & Walker, S. Fitting Linear Mixed-Effects Models Using lme4. J. Stat. Softw. 67, 1–48 (2015).

97. 97. Lenth, R. V. emmeans: Estimated Marginal Means, aka Least-Squares Means. R Package Version 180 https://CRAN.R-project.org/package=emmeans, (2022).

98. Allaire, J. M. et al. The Intestinal Epithelium: Central Coordinator of Mucosal Immunity. Trends Immunol. 39, 677–696 (2018).

99. Migone, T.-S. et al. TL1A Is a TNF-like Ligand for DR3 and TR6/DcR3 and Functions as a T Cell Costimulator. Immunity 16, 479–492 (2002).

100. Dheer, R. et al. Intestinal Epithelial Toll-Like Receptor 4 Signaling Affects Epithelial Function and Colonic Microbiota and Promotes a Risk for Transmissible Colitis. Infect. Immun. 84, 798–810 (2016).

101. Kim, S.-H. et al. Structural requirements of six naturally occurring isoforms of the IL-18 binding protein to inhibit IL-18. Proc. Natl. Acad. Sci. U. S. A. 97, 1190–1195 (2000).

102. Beumer, J. et al. Enteroendocrine cells switch hormone expression along the crypt-to- villus BMP signaling gradient. Nat. Cell Biol. 20, 909–916 (2018).

103. Roth, K. A., Kim, S. & Gordon, J. I. Immunocytochemical studies suggest two pathways for enteroendocrine cell differentiation in the colon. Am. J. Physiol. 263, G174–180 (1992).

104. Elmentaite, R. et al. Cells of the human intestinal tract mapped across space and time. Nature 597, 250–255 (2021).

105. Xie, Q. et al. The Role of Kisspeptin in the Control of the Hypothalamic-Pituitary- Gonadal Axis and Reproduction. Front. Endocrinol. 13, 925206 (2022).

106. Tomaro-Duchesneau, C. et al. Discovery of a bacterial peptide as a modulator of GLP-1 and metabolic disease. Sci. Rep. 10, 1–12 (2020).

107. Breton, J. et al. Gut Commensal E. coli Proteins Activate Host Satiety Pathways following Nutrient-Induced Bacterial Growth. Cell Metab. 23, 324–334 (2016).

108. Rabiei, S., Hedayati, M., Rashidkhani, B., Saadat, N. & Shakerhossini, R. The Effects of Synbiotic Supplementation on Body Mass Index, Metabolic and Inflammatory Biomarkers, and Appetite in Patients with Metabolic Syndrome: A Triple-Blind Randomized Controlled Trial. J. Diet. Suppl. 16, 294–306 (2019).

109. Ye, L. et al. Enteroendocrine cells sense bacterial tryptophan catabolites to activate enteric and vagal neuronal pathways. Cell Host Microbe 29, 179–196.e9 (2021).

110. De Vadder, F. et al. Microbiota-Generated Metabolites Promote Metabolic Benefits via Gut-Brain Neural Circuits. Cell 1–13 (2014) doi:10.1016/j.cell.2013.12.016.

111. Yano, J. M. et al. Indigenous Bacteria from the Gut Microbiota Regulate Host Serotonin Biosynthesis. Cell 161, 264–276 (2015).

112. Yaghoubfar, R. et al. The impact of *Akkermansia muciniphila* and its extracellular vesicles in the regulation of serotonergic gene expression in a small intestine of mice. Anaerobe 83, 102786 (2023).

113. Busslinger, G. A. et al. Human gastrointestinal epithelia of the esophagus, stomach, and duodenum resolved at single-cell resolution. Cell Rep. 34, 108819 (2021).

114. Long, A. et al. Famsin, a novel gut-secreted hormone, contributes to metabolic adaptations to fasting via binding to its receptor OLFR796. Cell Res. 33, 273–287 (2023).

115. Coll, A. P. et al. GDF15 mediates the effects of metformin on body weight and energy balance. Nature 578, 444–448 (2020).

116. Hu, X. et al. A gut-derived hormone regulates cholesterol metabolism. Cell 187, 1685–1700.e18 (2024).

117. Dhanvantari, S., Seidah, N. G. & Brubaker, P. L. Role of prohormone convertases in the tissue-specific processing of proglucagon. Mol. Endocrinol. 10, 342–355 (1996).

118. Bohórquez, D. V. et al. An Enteroendocrine Cell – Enteric Glia Connection Revealed by 3D Electron Microscopy. PLoS ONE 9, e89881 (2014).

119. Izzi-Engbeaya, C. & Dhillo, W. S. Gut hormones and reproduction. Ann. Endocrinol. 83, 254–257 (2022).

120. Uhlén, M. et al. Tissue-based map of the human proteome. Science 347, 1260419 (2015).

121. Kimura, I. et al. The gut microbiota suppresses insulin-mediated fat accumulation via the short-chain fatty acid receptor GPR43. Nat. Commun. 4, 1829 (2013).

122. Chambers, E. S. et al. Effects of targeted delivery of propionate to the human colon on appetite regulation, body weight maintenance and adiposity in overweight adults. Gut 64, 1744–1754 (2015).

123. Lund, M. L. et al. Enterochromaffin 5-HT cells – A major target for GLP-1 and gut microbial metabolites. Mol. Metab. 11, 70–83 (2018).

124. Chimerel, C. et al. Bacterial Metabolite Indole Modulates Incretin Secretion from Intestinal Enteroendocrine L Cells. Cell Rep. 9, 1202–1208 (2014).

125. Wang, Q. et al. Gut microbiota regulates postprandial GLP-1 response via ileal bile acid- TGR5 signaling. Gut Microbes 15, 2274124 (2023).

126. Ruan, W. et al. Enhancing responsiveness of human jejunal enteroids to host and microbial stimuli. J. Physiol. 598, 3085–3105 (2020).

